# Conformational rearrangements in 2^nd^ voltage sensor domain switch PIP_2_- and voltage-gating modes in two-pore channels

**DOI:** 10.1101/2022.06.14.494918

**Authors:** Takushi Shimomura, Kiichi Hirazawa, Yoshihiro Kubo

## Abstract

Two-pore channels (TPCs) are activated by PIP_2_ binding to domain I and/or by voltage-sensing in domain II (DII). Little is known about how these two stimuli are integrated and how each TPC subtype achieves its unique preference. Here, we show that the distinct conformations of DII-S4 in the voltage-sensor domain determine the two gating modes. DII-S4 takes an intermediate conformation, and forced stabilization in this conformation was found to give or maintain a high PIP_2_-dependence in primarily voltage-dependent TPC3 or in PIP_2_-gated and non-voltage-dependent TPC2, respectively. We also found in TPC2 that a tricyclic antidepressant desipramine induces the DII-S4 based voltage-dependence and that a flavonoid naringenin biases the mode preference from PIP_2_-gating to desipramine-induced voltage-gating. Taken together, our study on TPCs revealed an unprecedented mode-switching mechanism involving conformational changes in DII-S4. This will pave the way for drug development by targeting specific gating modes of TPCs.

**Significance statement:** Membrane voltage and PIP_2_ are different types of signals on endosomal and lysosomal membranes. The two signals are integrated into two-pore channels (TPCs) whose two repeating domains, DI and DII, play roles in PIP_2_ binding and voltage sensing, respectively. We showed that the conformation of the S4 helix in DII determines the voltage-dependent or PIP_2_-dependent gating mode, which explains the different preferences of the two signals between TPC subtypes. The preference for these two gating modes can be changed by a flavonoid, naringenin. Our findings on the molecular mechanism of the two gating modes in TPCs provide a clue to the understanding and pharmacological manipulation of the signaling by PIP_2_ and voltage in intracellular organelles.

## Introduction

Two-pore channels (TPCs) are members of of voltage-gated cation channel superfamily (1–3). TPCs play a major role in nicotinic acid adenine dinucleotide phosphate-dependent Ca^2+^ release from intracellular organelles such as lysosomes, endosomes, and cortical granules, which serve as Ca^2+^ stores that are different from the endoplasmic reticulum (4, 5). Consequently, TPCs are related to diverse physiological functions, including angiogenesis (6) and hair color determination (7), as well as pathophysiologies such as fatty liver disease (8) and Parkinson’s disease (9). Owing to the critical roles of TPCs in intracellular organelles, their pharmacological inhibition and gene knock-out have been known to prevent the intracellular entry of the Ebola virus (10). Similarly, severe acute respiratory syndrome coronavirus 2 (SARS-CoV-2), which has emerged as the cause of the COVID-19 pandemic, invades cells through angiotensin-converting enzyme 2-mediated endocytosis involving endosomes and lysosomes (11–13). Consistent with this notion, some TPC2 inhibitors, such as tetrandrine and naringenin, suppressed SARS-CoV-2 infection in *in vitro* assays (14–17).

TPCs are composed of two homologous domains (DI and DII), each of which contains six transmembrane helices that are the functional units of voltage-gated cation channels (1, 18, 19) and function as dimers (20, 21). The former four helices form a voltage sensor domain (VSD) and the latter two helices comprise a pore domain (PD). In VSDs, positively charged residues in helix S4 are generally important for voltage sensing (22–24). In the case of TPCs, while both DI-S4 and DII-S4 have positively charged residues, voltage sensitivity is primarily governed by DII-S4, and not by DI-S4 (21, 25, 26). Electrophysiological recordings have shown that TPCs are also activated by phosphatidylinositol bisphosphate (PIP_2_), PI(3, 5)P_2_, and/or PI(3, 4)P_2_ and selectively allow Na^+^ permeation (27–31). Several cryo-EM structures of TPCs have shown that PIP_2_ binds to a region composed of a DI-S4/S5 linker and VSD1 (25, 26, 32). Therefore, TPCs are a unique type of voltage-gated ion channel, in which each domain is responsible for ligand binding and voltage sensing.

It is still unclear how PIP_2_ binding and voltage sensing is integrated in TPCs, mainly owing to the lack of functional studies employing electrophysiological analysis, in contrast to the many existing structural reports. Notably, structures with apo- and PIP_2_- bound states have been previously reported (26, 32), where PIP_2_-induced structural rearrangements around the activation gate were mainly discussed in terms of the gating mechanism. However, our previous functional study of TPC3 from *Xenopus tropicalis* (XtTPC3) showed that PIP_2_ binding in DI affects the movement of distantly located DII- S4 (30), suggesting that the integration mechanisms of PIP_2_ and voltage in TPC3 are global and complex. In addition, TPC2, which is believed to be non-voltage-dependent, was recently reported to elicit a voltage-dependent current in the presence of a series of chemical compounds, named Lysosomal Na^+^ channel Voltage-dependent Activators (LyNaVAs) (33). Furthermore, another type of TPC2 agonist has been shown to change cation selectivity (34). TPC2 and perhaps other TPC subtypes have more complex integration mechanisms for PIP_2_ and voltage than expected. Moreover, it remains unclear how different preferences and sensitivities to the respective stimuli are achieved in each TPC subtype.

To explore the details of the activation mechanism of PIP_2_ and voltage in the TPCs, we first focused on XtTPC3. Voltage clamp fluorometry (VCF) analysis of XtTPC3 revealed that DII-S4 has an intermediate state that opens in a strongly PIP_2_-dependent manner. A mutation that stabilizes this intermediate state transforms XtTPC3, which is predominantly voltage-gated, into a strong PIP_2_-gated channel type, such as TPC2. Next, we focused on human TPC2 (HsTPC2) and revealed similarities in the PIP_2_- and voltage-gating modes in TPC subtypes with different mechanisms. Furthermore, with the presence of the two modes in mind, we found that naringenin is not a simple TPC2 inhibitor rather it acts to bias the equilibrium between the two modes. These results reveal a unique regulatory mechanism that is likely to be common among TPC subtypes, where DII-S4 conformations define PIP_2_-gating and voltage gating. These new findings will form the basis for a new approach to drug development against SARS-CoV-2 and other viruses.

## Results

### Two steps of movements of DII-S4 in XtTPC3 revealed by VCF analysis

VCF is a method used to track local structural changes using a fluorophore introduced to a specific amino acid. When a fluorophore is attached to the top of the S4 helix of typical voltage-gated ion channels, such as a *Shaker* channel, the fluorescent changes reflect, in most cases, the movement corresponding to the transfer of gating charges that precede the opening of the activation gate. Therefore, voltage-dependent changes in fluorescence intensity are generally detected at more hyperpolarized voltages than currents, and match the voltage dependence of the gating charge transfer (Fig. 1a) (35). To analyze the movement of DII-S4, we performed VCF analysis using XtTPC3 with a fluorophore attached to a cysteine introduced at Gln507, which is located at the extracellular end of DII-S4 (Fig. 1b). Current and fluorescence were simultaneously measured in the presence of PI(3, 4)P_2_, which potentiates the voltage dependence of XtTPC3 (31). PI(3, 4)P_2_ levels were increased by co-expression of human INPP5D (HsIP5D), a PI(3, 4)P_2_ producing enzyme, and insulin treatment (36). We observed changes in fluorescence intensity that were made of two distinct components (Fig. 1c–e). One component with a gradual voltage-dependent decrease (F1) was observed at more hyperpolarized potentials than for those where channel opening occurred, whereas the other (F2) showed a relatively steep increase at more depolarized potentials. As two components were observed in the distinct range of the membrane voltages, at a constant +90 mV step pulse, a rapid fluorescence decrease corresponding to F1 and a subsequent gradual increase derived from F2 could be resolved (Fig. 1d). These two types of fluorescence intensity changes with different properties indicate that DII-S4 has an intermediate conformation during the transition from a resting state to a fully activated state. Our previous recordings showed that the major gating charge movement in XtTPC3 occurred in the range of membrane potentials in which F2 was observed (Fig. 1e) (30). Taken together, the DII-S4 movement in XtTPC3 is thought to be quite different from typical S4 movements (Fig. 1a, e).

**Figure 1.**
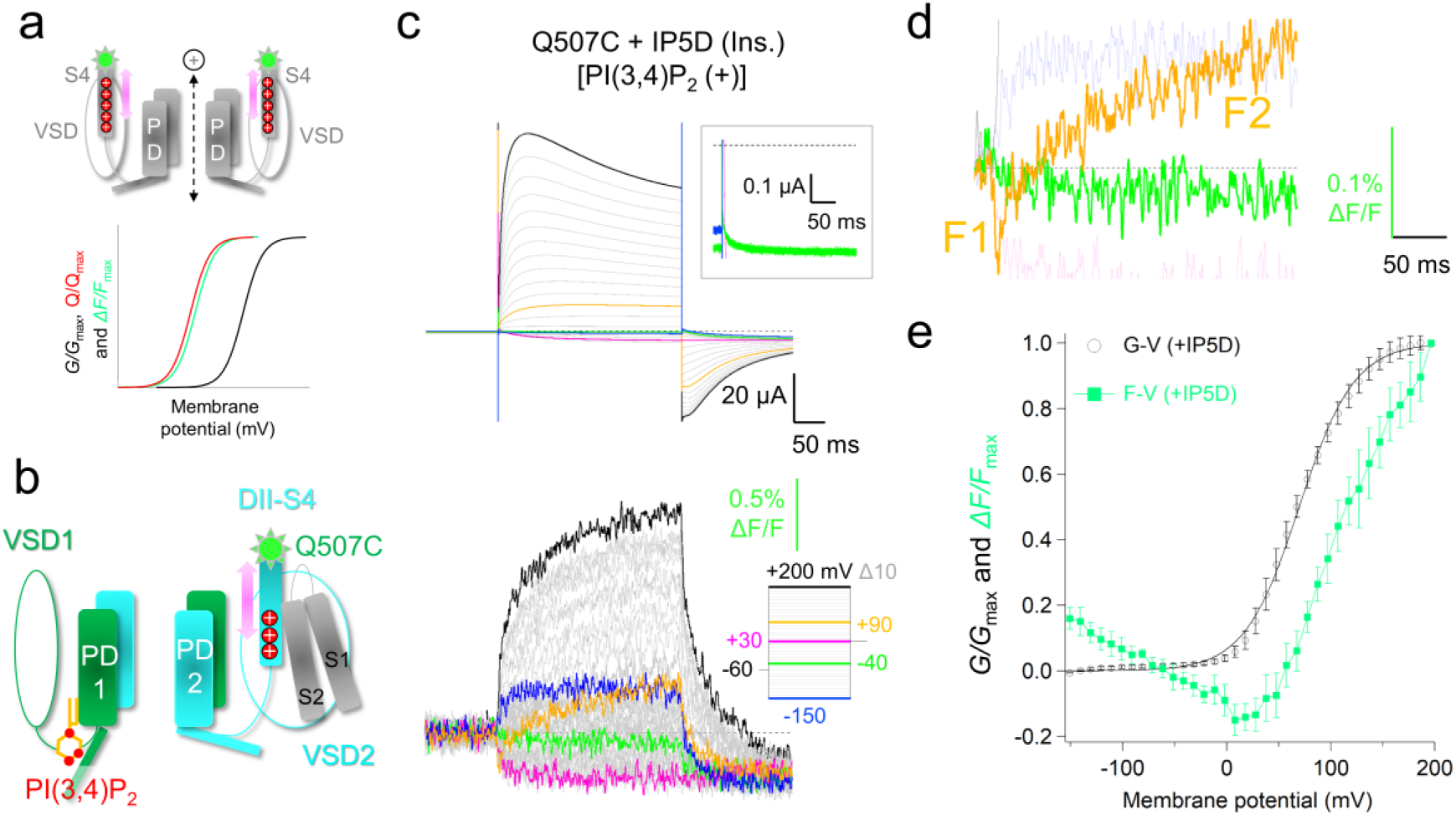
Two components of fluorescent changes in the VCF measurement of XtTPC3. (a) A schematic drawing for the typical voltage-gated ion channel with the S4 helix labelled by a fluorophore (top) and an example of its VCF measurement (bottom). Two VSDs and their linkers to the PDs were omitted for clarity. *G/G*_max_, *Q/Q*_max_ and *ΔF/F*_max_ indicate the voltage dependence of normalized conductance, gating charge and changes in fluorescence intensity, respectively. (b) A cartoon for the molecular structure of XtTPC3. VSD1 and VSD2 from one subunit and their linkers to the PDs were omitted for clarity. DII-S1 and DII-S2 (grey), and DII-S4 (aqua) are shown on a highlighted VSD2. Asn507 (green light spot) and 3 positively charged residues (red) are also depicted. (c) Representative current recordings (upper) and fluorescence recordings (lower) for Q507C attached with Alexa-488. The expanded view of the current trace elicited by a -40 mV pulse from a holding potential at -60 mV is also shown (inset). The step pulses to elicit the current and fluorescence are shown on the right side of the fluorescence in matched colors of the corresponding traces. (d) The expanded view of the fluorescence traces in (c), in which the region onset of the current deviation by the step pulses to -40 mV and +90 mV were shown. The color codes are the same as in (c). (e) Normalized ΔF-V (green, n = 7) and G-V relationships in Q507C (black, V_1/2_ = 69.5 ± 0.84, n = 6).

### The introduction of the electrostatic interaction between Glu447 and Phe514 confers an additional activation component that is highly sensitive to PI(3, 4)P_2_

In contrast to the two clear components measured in the VCF analysis, only a single component in the conductance-voltage (G-V) relationship was observed at the depolarized membrane potentials, suggesting that the intermediate state does not lead to a sufficient level of channel opening in WT XtTPC3 (Fig. 2a). A mutation in DII-S4 that uncovers this hidden state was searched for to investigate the intermediate state in detail. Substitution of Phe514 with arginine resulted in channel opening, even at -60 mV (Fig. 2a inset) and a clear two-step voltage dependence in the current in the presence of PI(3, 4)P_2_ (Fig. 2b). Of these two steps, the one on the high-voltage (HV) side occurred in a range similar to the voltage dependence in WT XtTPC3, while the other occurred on the very low voltage (LV) side with a less steep slope. Notably, the LV component in F514R appeared in a similar membrane potential range, where the F1 component was observed in the VCF of Q507C (Fig. 2b).

**Figure 2.**
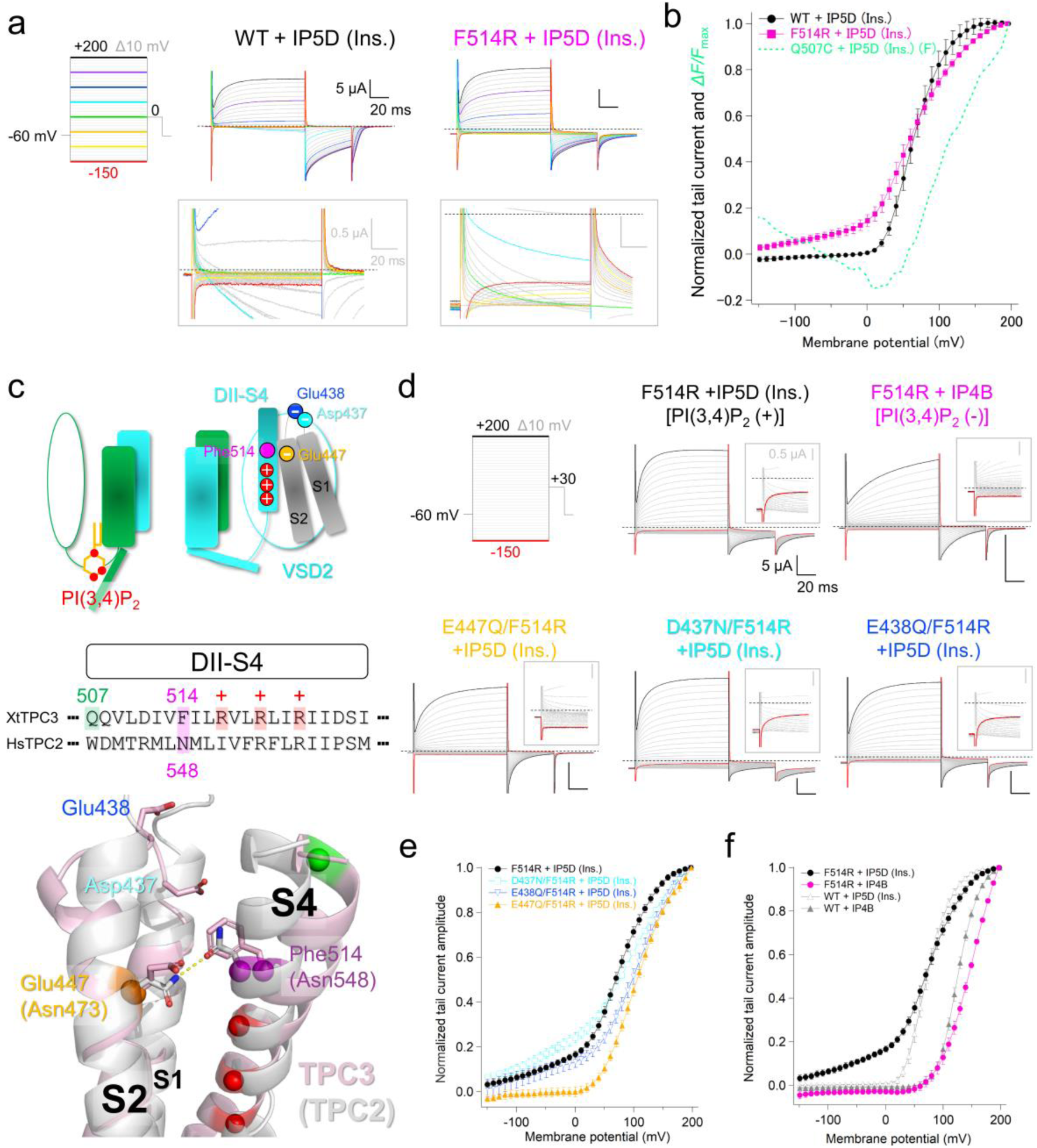
Two steps of voltage-dependent activation in F514R. (a) Step pulses to elicit each current that are colored to indicate the corresponding current traces (left), and the representative current recordings of WT (middle) and F514R (right). The expanded views to highlight the currents elicited by hyperpolarized step pulses are also shown (inset). (b) Normalized tail current amplitude of WT (black, n = 5) and F514R (magenta, n = 4) at each membrane potential. The same plot for ΔF-V of Q507C as in Fig. 1e is shown (dotted green line) for comparison. (c) A cartoon for the molecular structure of XtTPC3 (top), amino acid sequence alignments of DII-S4 of XtTPC3 and HsTPC2 (middle) and the aligned structures of XtTPC3 and HsTPC2 (bottom). Each residue is depicted in matched colors in all three panels. DII-S3 was omitted for clarity. The spheres indicate the C*α* positions of each colored residue. The yellow dotted line indicates the hydrogen bond between Asn473 and Asn548 in HsTPC2. (d) Representative current recordings in each mutant, elicited by the step pulses from -150 mV to +200 mV (left and top). The expanded views of the current traces around the onset of current deviation are also shown to check the conductance at hyperpolarized potentials in each mutant (inset). (e) Normalized tail current amplitude in a series of mutants at each membrane potential (n = 4-5). (f) Normalized tail current amplitude in F514R or WT, co-expressed with HsIP4B or HsIP5D respectively, at each membrane potential (n = 4).

We hypothesized that the LV component is caused by an electrostatic interaction formed between the introduced F514R and a negatively charged residue endogenous to VSD2, such as Asp437, Glu438, and Glu447 (Fig. 2c). Although additional D437N or E438Q mutations to F514R still showed the LV component with an apparent shift in the HV component, the LV component was lost in E447Q/F514R in the presence of PI(3, 4)P_2_ (Fig. 2d, e). The XtTPC3 structural model, based on the cryo-EM structure of zebrafish TPC3 that is proposed to be in a deactivated state (37), shows that Phe514 is located near Glu447 in this conformation and is likely to approach Asp437 and Glu438 when DII-S4 is fully activated (Fig. 2c). Taken together, these results show that the conductance of the LV component appears when F514R is locked close to Glu447 in a deactivated-like VSD2 conformation, which is a distinct mechanism from the typical voltage-dependent opening that is accompanied by large S4 movement.

Our previous studies on XtTPC3 have shown that PI(3, 4)P_2_ binding to DI enhances the voltage-dependent movement of DII-S4, through the inter-domain interaction between DI-S6 and DII-S6 (30, 31). The recordings in this study were carried out in the presence of PI(3, 4)P_2_ by co-expression of HsIP5D and insulin treatment at a sufficiently high concentration to potentiate the voltage dependence of XtTPC3. In contrast, co-expression of WT XtTPC3 with human INPP4B (HsIP4B), which specifically degrades PI(3, 4)P_2_, significantly shifted the G-V relationship toward the depolarized direction (Fig. 2f and Supplementary Fig. 1). In F514R, HsIP4B co-expression resulted in a shift of the HV component toward the depolarized direction as in WT XtTPC3, however, it also showed a clear loss of the LV component, instead of a shift (Fig. 2d, f), suggesting that the LV component not only depends on the VSD conformation in DII but also on the PI(3, 4)P_2_ binding to DI.

### The disulfide-bond formation between Glu447 and Phe514 opens the channels in a PI(3, 4)P_2_-dependent manner

To further demonstrate the channel opening in the LV state, an oxidant that forms disulfide bonds, copper-phenanthroline (Cu-Phen), was applied to E447C/F514C and E438C/F514C. In the presence of PI(3, 4)P_2_, Cu-Phen treatment of both E447C/F514C and E438C/F514C increased the channel current (Fig, 3a, b). No clear changes in currents were observed following the Cu-Phen application to the single cysteine mutants or following the application of Cu^2+^ and phenanthroline alone to E447C/F514C (Supplementary Fig. 2). Because the currents increased over the entire range of the membrane potentials, including the hyperpolarized side, the disulfide-bond formations are likely to make the channels constitutively open (Fig. 3c, d). The remaining and unchanged voltage dependence after Cu-Phen treatment could be explained as that the currents are composed not only of the crosslinked and constitutively open fraction but also of the un-crosslinked fraction that opens in a voltage dependent manner as before the Cu-Phen application (Supplementary Table 1). These results indicate that channel opening was achieved in both conformations of DII-S4, where Phe514 was located in the proximity of Glu438 or Glu447, in the presence of PI(3, 4)P_2_. However, in the absence of PI(3, 4)P_2_, while currents after Cu-Phen treatment in E438C/F514C on the hyperpolarized side were observed, as in the presence of PI(3, 4)P_2_, those in E447C/F514C were significantly attenuated (Fig. 3c-e). These results, consistent with those of F514R, show that the conductance that occurs when Phe514 is close to Glu447 depends strongly on PI(3, 4)P_2_, unlike the typical voltage-dependent opening in which Phe514 on DII-S4 is located close to Glu438.

**Figure 3.**
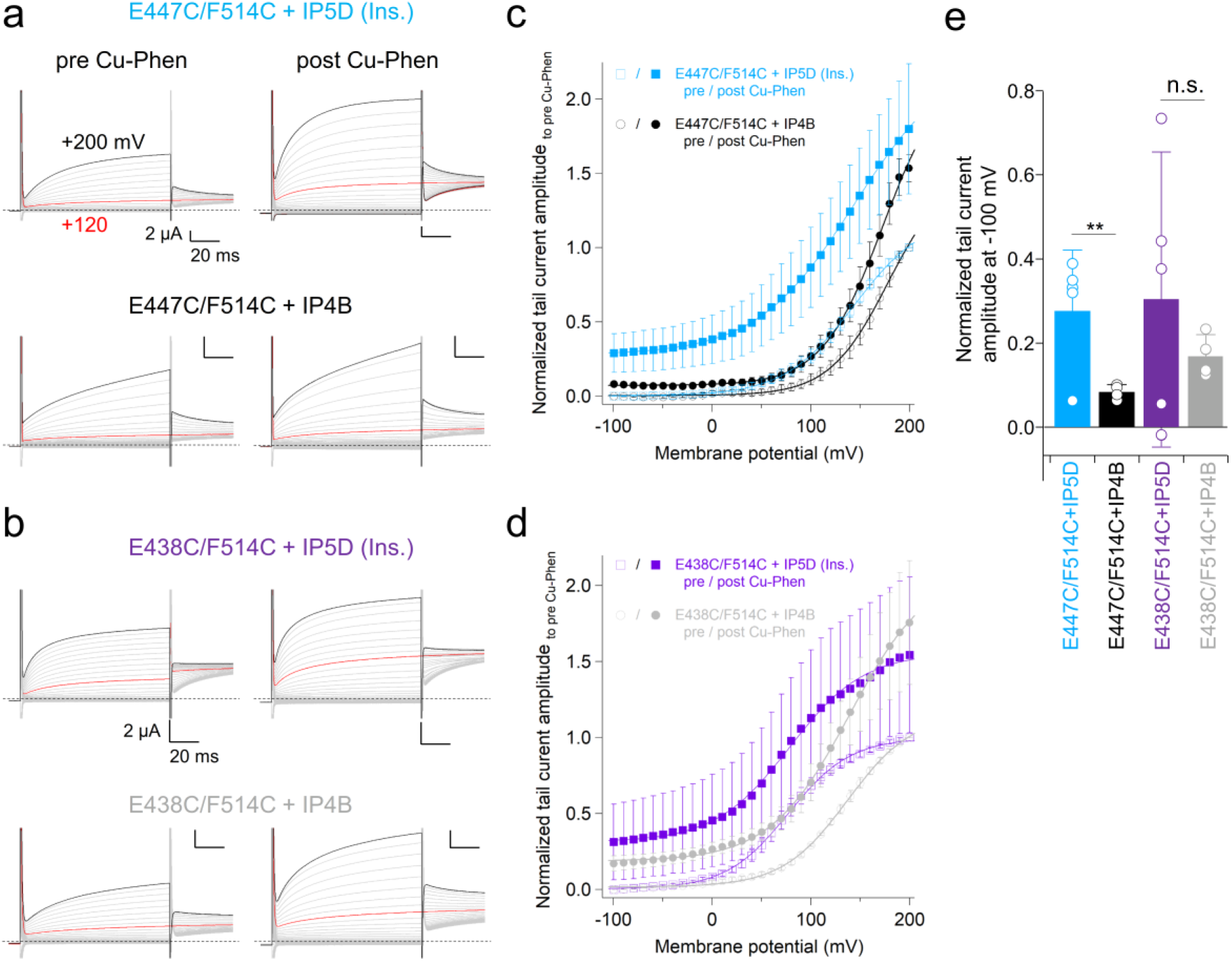
Disulfide bond formation in E447C/F514C and E438C/F514C, co- expressed with HsIP5D or HsIP4WB. (a, b) Representative current recordings of E447C/F514C (a) and E438C/F514C (b), co- expressed with HsIP5D and treated with insulin (top), or co-expressed with HsIP4B (bottom), respectively (n = 4-7). The recordings before (left) and after (right) Cu-Phen treatment are shown. The recordings were performed in the 20 mM Na^+^ solution to avoid too large an inward current through the constitutively open channels. (c, d) Normalized tail current amplitude at each membrane potential of E447C/F514C (c) and E438C/F514C (d), before and after Cu-Phen treatment. In all the combinations, the tail current amplitude was normalized to that before Cu-Phen treatment. The color codes for each combination are the same as used in (a) and (b). The open and filled symbols indicate the amplitude before and after Cu-Phen treatment, respectively. (e) Normalized tail current amplitudes at -100 mV after Cu-Phen treatment. Statistical significance was examined by unpaired t-test. ** and n.s. mean *p* < 0.01 and p > 0.05, respectively.

### The F1 component reflects the movement of DII-S4 to a conformation in which the channel opens with strong PI(3,4)P_2_-dependence

To confirm the correlation between the F1 component observed in the VCF of Q507C and the LV component of the F514R current, we performed a VCF analysis of the mutant that stabilizes the LV conformation. E447R/F514E was designed, with reference to the strategy used by Taylor et al. in the analysis of the KCNQ1 channel (38), to strongly stabilize the LV conformation by inducing two electrostatic repulsions and one interaction (Fig. 4a). E447R/F514E showed currents with a very gentle voltage dependence over the entire range of the measured membrane potentials (Fig. 4b, c). Notably, when HsIP4B was co-expressed to suppress the PI(3,4)P_2_ level, E447R/F514E produced almost no current over the entire range of membrane potentials. This is in sharp contrast to the WT current and the HV component of F514R, where channel currents remained even in the absence of PI(3,4)P_2_ (Fig. 2f, 4c), indicating that the E447R/F514E current consisted only of the LV component. VCF analysis of E447R/Q507C/F514E labeled with the fluorophore showed that the F1 was observed, as in the case of Q507C, while the F2 was almost completely lost (Fig. 4d, e). These results demonstrate that the conductance of the LV component, which has a weak voltage dependence and a strong PI(3,4)P_2_-dependence, is caused by the DII-S4 movement that corresponds to F1. Taken together with the previous results, it was shown that the LV component reflects channel opening in a conformation with a VSD2/activation gate in the intermediate/open (I/O) state, and that the HV component, corresponding to the F2 components, reflects opening in a fully activated/open (A/O) state.

**Figure 4.**
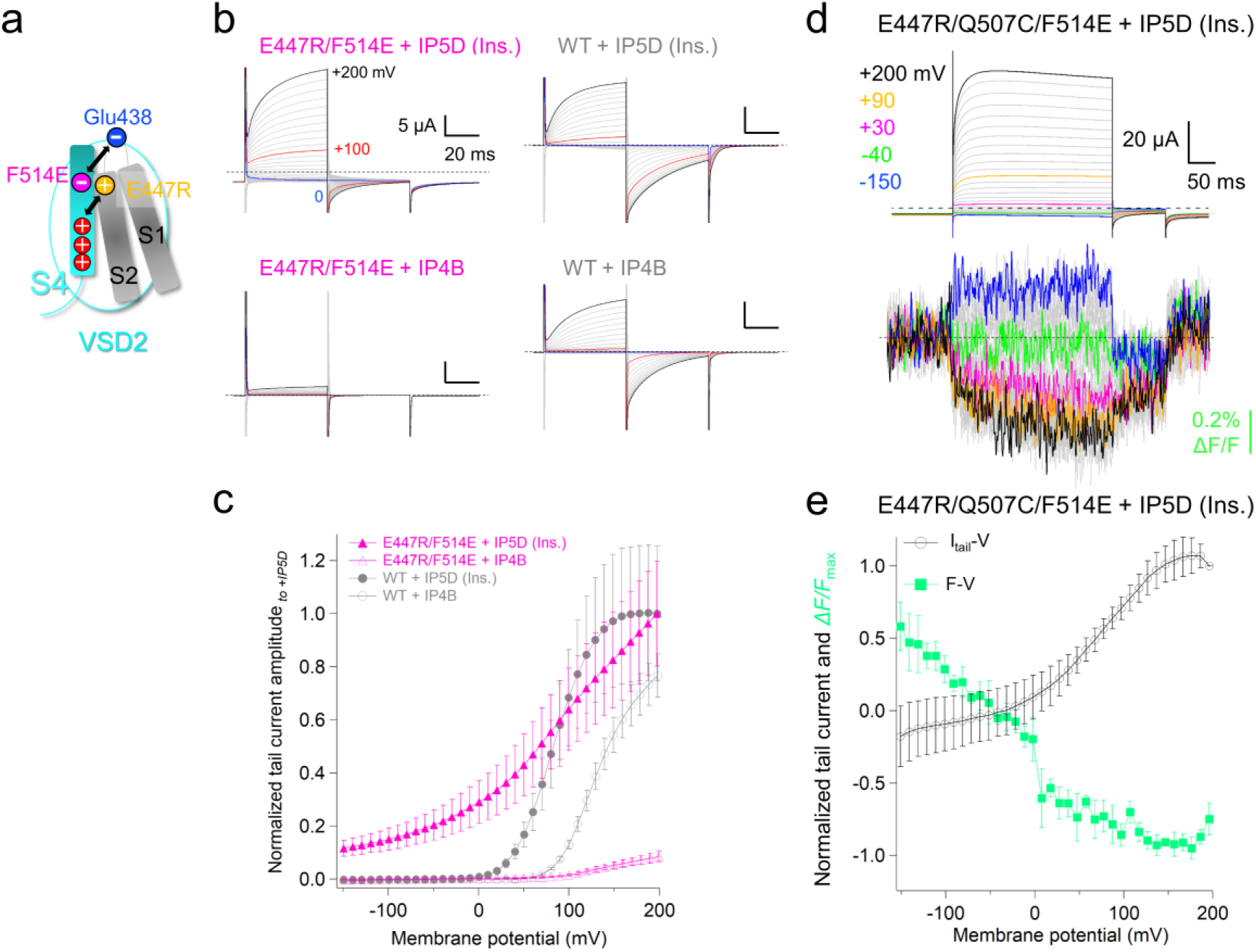
Current and fluorescence measurements of the E447R/F514E-type mutants. (a) A cartoon for the VSD2 domain to show the strategy to stabilize the LV component. The color codes are the same as in Fig. 2c. Arrowheads indicate electrostatic repulsion. (b) Representative current recordings of E447R/F514E (left) and WT (right), co-expressed with HsIP5D and treated with insulin (top), or HsIP4B (bottom), respectively. Currents are elicited by the step pulses from -150 mV to +200 mV from a holding potential of -60 mV, followed by a 0 mV pulse for 50 ms to elicit the tail currents. (c) Normalized tail current amplitudes of E447R/F514E (magenta) and WT (grey), co-expressed with HsIP5D and treated with insulin (filled), or HsIP4B (open), respectively (n = 5). In both WT and E447R/F514E, the tail current amplitude was normalized to the averaged amplitude of the tail currents in the HsIP5D co-expression and insulin treatment condition. (d) Representative current recordings (upper) and fluorescence recordings (lower) for E447R/Q5070C/F514E co-expressed with HsIP5D and treated by insulin, and labelled by Alexa-488. Currents and fluorescence are elicited by the step pulses for 300 ms, ranging from -150 mV to +200 mV, from a holding potential of -60 mV, followed by a 0 mV pulse for 100 ms to elicit the tail currents. The recordings were performed in the ND96-based solution containing 20 mM NaCl to decrease the currents at the holding potential. (e) Normalized ΔF-V (green; n = 4) and normalized tail current amplitude (black, n = 4)at each membrane potential of E447R/Q507C/F514E co-expressed with HsIP5D and treated by insulin.

### The effects of PI(3,4)P_2_ on the F1 and F2 components

Because the LV component in F514R, which corresponds to the F1 component and I/O state, has a strong PI(3,4)P_2_ dependence, we performed a VCF with Q507C in the absence of PI(3,4)P_2_ and compared the results with those in the presence of PI(3,4)P_2_. In the VCF measurement of Q507C co-expressed with HsIP4B, a similar F1 component was observed (Fig. 5a, b). In contrast to the measurement in the presence of PI(3,4)P_2_, where F2 followed G-V, the voltage dependence of F2 almost overlapped with G-V in the absence of PI(3,4)P_2_ (Fig. 1e, 5b). To further investigate the coincidence of the current and F2, the time courses of the simultaneous recordings were compared in step pulses at the membrane potentials where F2 could be observed (Fig. 5c and Supplementary Fig. 3). At each membrane potential, while the time constants for current were significantly smaller than those for F2 in the presence of PI(3,4)P_2_, they were not significantly different from each other in its absence (Fig. 5c, d), indicating that the channel opening was already achieved before the F2 movement when PI(3,4)P_2_ was bound, but concurrently occurred when unbound.

**Figure 5.**
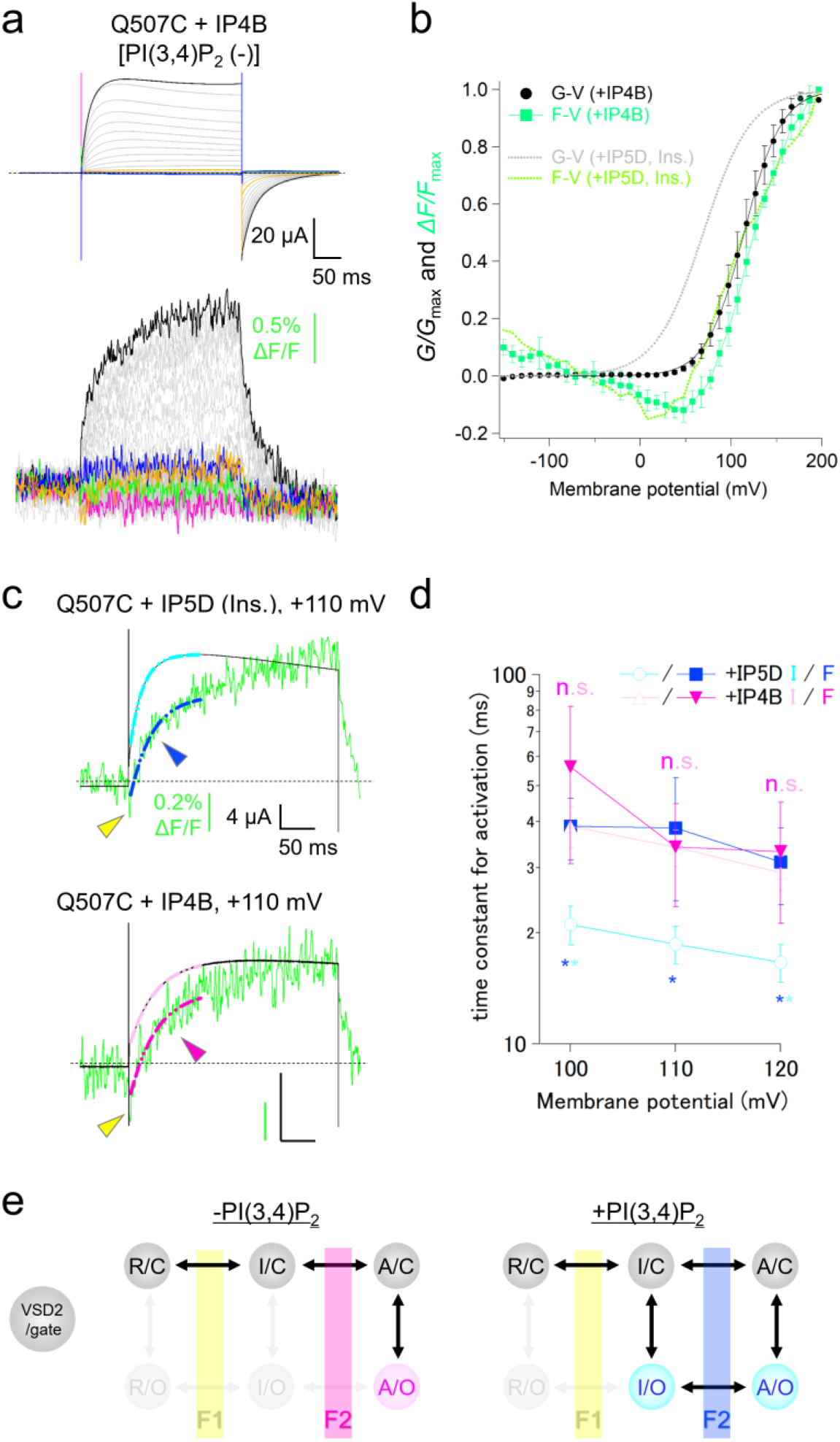
VCF analysis of the PI(3,4)P_2_-dependence in Q507C. (a) Representative recordings for current (upper) and fluorescence (lower) of Q507C co- expressed with HsIP4B labelled by Alexa-488. Currents and fluorescence are elicited by the same pulse protocol shown in Fig. 1c. (b) Normalized ΔF-V (pale green; n = 8) and G-V relationships in Q507C co-expressed with HsIP4B (black, V_1/2_ = 114 ± 0.46, n = 8). The ΔF-V (dotted yellow-green line) and the fitted G-V curve (dotted grey line) of Q507C co-expressed with HsIP5D shown in Fig. 1e are also plotted for comparison. (c) Superimposed recordings of current and fluorescence of Q507C co-expressed with HsIP5D and treated with insulin (top) and HsIP4B (bottom). In both cases, the fluorescence recordings (blue and magenta, respectively) and current recordings (aqua and pink, respectively) were fitted by a single exponential function. The yellow arrowheads indicate the F1 components, while the blue and magenta arrowheads indicate the F2 components. (d) The activation time constants are obtained in (c) at each membrane potential (n = 6-7). The color codes match those used in (c). (e) Schematic diagrams for state transitions of XtTPC3 in the absence (left) and presence (right) of PI(3, 4)P_2_. R, I, and A in the spheres indicate the VSD2 conformations in resting, intermediate, and activated states, respectively. C and O indicate the conformations of the activation gate in the closed and open states, respectively. The regions colored and denoted as F1 or F2 indicate the transitions corresponding to each fluorescent intensity change. The faint spheres and arrowheads indicate their low state occupancy and low rates of state transitions, respectively. Paired t-test was used to examine statistical significance. **, * and n.s. mean *p* < 0.01, *p* < 0.05, and p > 0.05, respectively.

The results of E447R/F514E, which lacked the F2 component and consisted only of F1, demonstrated that the intermediate state, to which F1 reflects the transition, can lead to channel opening in the presence of PI(3,4)P_2_ (Fig. 4). Therefore, it is natural to assume that even in WT XtTPC3, the I/O state occurs in the presence of PI(3,4)P_2_ and contributes to channel opening before F2 (Fig. 5e). Consistent with this assumption, a small but clear current in the WT channel was observed at the hyperpolarized potential where the F1 component is dominant, in the measurement with a high PI(3,4)P_2_ level (Fig. 1c inset). Because no channel opening was observed until F2 in the absence of PI(3,4)P_2_ (Fig. 5c), the transition into the I/O state is likely to be quite limited. Instead, the opening in the A/O through the activated/closed (A/C) state could occur concurrently with F2 which reflects the transition from I/C to A/C (Fig. 5e). Assuming this scheme, the G-V relationship of WT XtTPC3 in the presence of PI(3,4)P_2_ contains two components: one in a hyperpolarized range derived from the I/O state, which is small owing to its low state occupancy, and the other in the depolarized range from the A/O state, with a high open probability that is facilitated by the gating charge transfer corresponding to F2 (Fig. 5e). Thus, a depolarizing stimulus to WT XtTPC3 in the presence of PI(3,4)P_2_ is likely to trigger the current from the I/O state with a low open probability, followed by a slower A/O-derived current with a high open probability (Fig. 5c). These observations suggest that TPC3 intrinsically has a unique I/O state that is strongly regulated by PI(3,4)P_2_.

### The correspondence of the two gating modes in XtTPC3 to the LyNaVA- and PI(3, 5)P_2_-dependent gating modes in HsTPC2

The results so far show that XtTPC3 is intrinsically designed to open in two distinct DII-S4 conformations: one is the A/O state with DII-S4 fully activated, and the other is the I/O state with less voltage-dependence and strong PI(3, 4)P_2_-dependence. To investigate whether these two gating mechanisms are shared in the TPC family, we next focused on TPC2 which has distinct features from those of TPC3.

It is known that TPC2 has a LyNaVA-mediated voltage dependence. Because the molecular mechanisms of LyNaVA-induced currents remain elusive (33), we examined whether this unusual voltage dependence is regulated by DI- or DII-S4. A ΔN37 mutant of human TPC2 (HsTPC2), in which those N-terminal region critical for lysosomal retention was deleted (39, 40), was used to express HsTPC2 on the plasma membrane. HsTPC2 ΔN37, hereafter referred to as WT, showed a current in response to the intracellular microinjection of PI(3, 5)P_2_ (Fig. 6a and Supplementary Fig. 4a, b). Extracellular perfusion with desipramine, a type of LyNaVA, also induced currents in the absence of PI(3, 5)P_2_ (Fig. 6a). While PI(3, 5)P_2_-evoked currents showed no voltage-dependence, desipramine-induced currents showed a clear voltage-dependence and inward rectification (V_1/2_ = -45.0 ± 4.2, n = 6) (Fig. 6a, b, and Supplementary Fig. 4c), demonstrating that desipramine-induced HsTPC2 currents were reproduced as previously reported, even in the oocyte expression system (33). Neutralizing mutations of arginine residues in DI-S4, such as R185Q and R188Q, did not significantly change the V_1/2_ values of desipramine-induced currents. However, significant shifts toward the depolarized direction were observed in the R554Q and R557Q mutants of DII-S4 (Fig. 6a, b, and Supplementary Fig. 4d, e, Supplementary Table 1). The I551R mutant, which adds a positive charge at an upper position in DII-S4 (Fig. 2c), did not show a current with voltage dependence under basal conditions. In the presence of desipramine, it showed a current with the G-V shifted toward a positive direction compared to WT, possibly due to the perturbation of the optimal voltage-dependent movement of DII-S4 by introducing an extra positive charge. These results show, for the first time, that LyNaVA-induced voltage-dependent currents in HsTPC2 depend on DII-S4.

**Figure 6.**
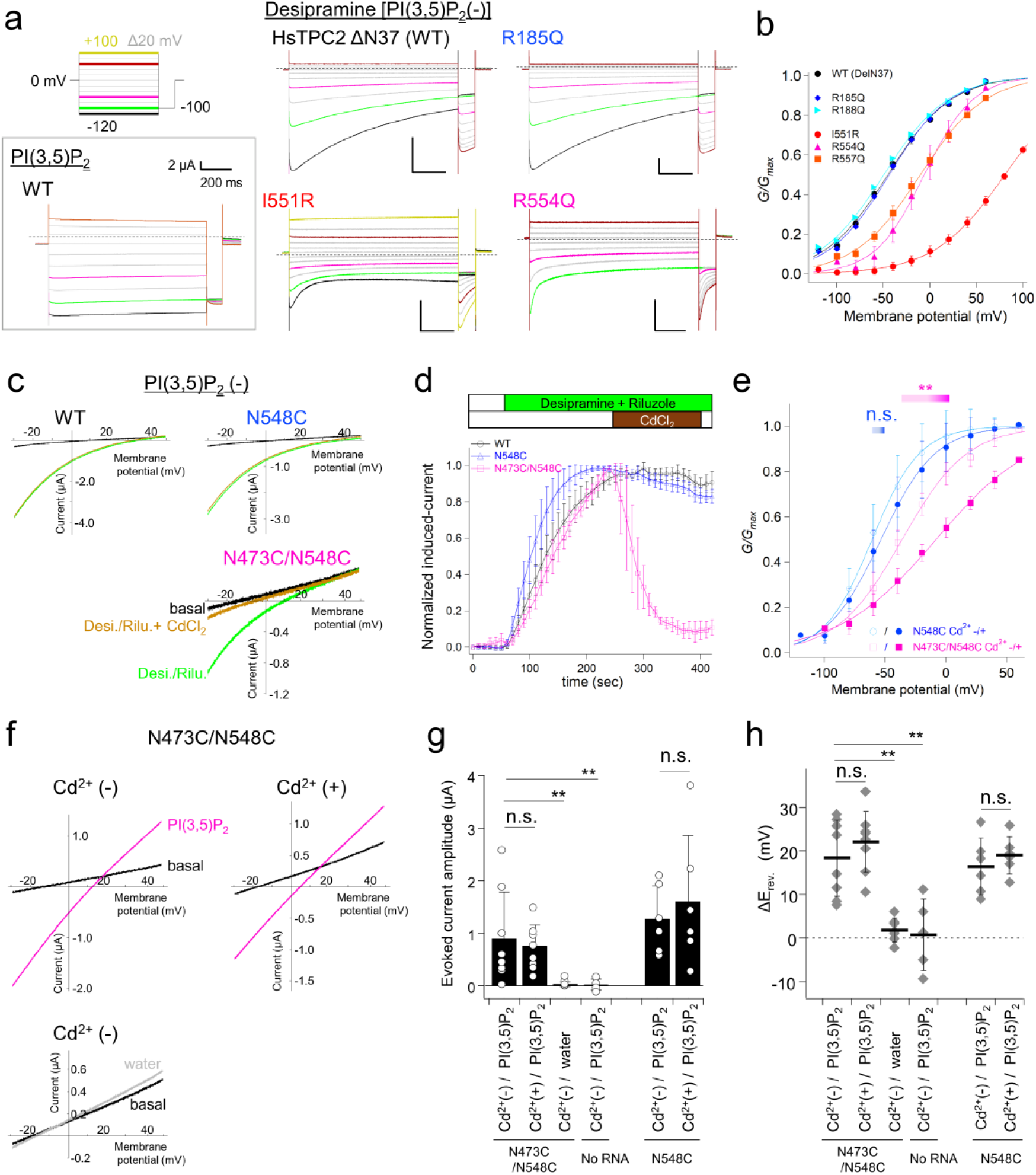
Comparative analysis of desipramine-induced and PI(3,5)P_2_-evoked currents in HsTPC2. (a) Step pulses to elicit the currents that are colored to indicate the corresponding current recordings (top left) and the representative current recordings of HsTPC2 WT and mutants. Desipramine-induced currents and PI(3, 5)P_2_-injection evoked current (grey box) in WT and the three mutants, with ΔN37 as common background are shown. (b) G-V relationships of WT and the RQ mutants obtained from the tail current at -100 mV in (a) (n = 4-6). (c) Representative recordings of desipramine-induced currents in each HsTPC2 construct. Currents were evoked by ramp pulses ranging from -30 mV to +50 mV. TPC2 currents were induced by the co-application of 1 mM desipramine and 100 µM riluzole, followed by the addition of 100 µM CdCl_2_. (d) Time courses of the change in current amplitude of HsTPC2 in response to the application of desipramine/riluzole and then CdCl_2_ (n = 4). Desipramine/riluzole-induced currents were normalized to the maximum induced currents. (e) G-V relationships of N548C and N473C/N548C with (filled) or without (open) 100 µM CdCl_2_ in the bath solution (n = 5-7). (f) Representative currents recordings of N473C/N548C mutant before (black) and after the injection of 5 mM PI(3, 5)P_2_ (magenta) or water (grey) in the presence or absence of 100 µM CdCl_2_ in the bath solutions. Currents were evoked by ramp pulses ranging from -30 mV to +50 mV. (g and h) The plots of the current amplitude (g) and the shift of the reversal potential (h) evoked by PI(3, 5)P_2_ or water injection into the oocyte expressing TPC2 mutants or no exogenous proteins (n = 5-8). Statistical significance was examined by unpaired t-test or One-way ANOVA followed by Dunnett’s test in (e) and (g, h), respectively. ** and n.s. mean *p* < 0.01 and p > 0.05, respectively.

It is noteworthy that the cryo-EM structures of HsTPC2 showed in both its apo- and PI(3, 5)P_2_ bound-states that Asn473 and Asn548, which correspond to Glu447 and Phe514 of XtTPC3, are located close enough to form a hydrogen bond (Fig. 2c). Assuming that the mechanisms for the PIP_2_- and voltage-gating modes in TPC2 are similar to those in TPC3, further locking of this conformation would perturb the desipramine-induced voltage-dependent opening mechanism due to the restricted movement of DII-S4, but still allow the PI(3, 5)P_2_-dependent mechanism. N473C/N548C, aimed at a Cd^2+^ coordination- induced bridge between the two residues, produced only a very small desipramine-induced current. To measure the time course of the effect of Cd^2+^ with a larger current amplitude, we co-perfused riluzole, another type of agonist different from LyNaVAs (33), and desipramine, to N473C/N548C (Fig. 6c). Further perfusion with Cd^2+^ strongly inhibited the desipramine/riluzole-induced current (Fig. 6c, d). Under the same conditions, the N473C current was undetectable, and no Cd^2+^ effects were observed in N548C. The application of desipramine alone to N473C/N548C also evoked small currents that were similarly attenuated by Cd^2+^ perfusion (Supplementary Fig. 5a, b). In addition, a Cd^2+^- induced shift in the G-V relationship of the desipramine-induced current toward the depolarized direction was also confirmed in N473C/N548C (Fig. 6e, Supplementary Fig. 5c, Supplementary Table 1). In contrast, the presence or absence of Cd^2+^ caused no significant differences in PI(3, 5)P_2_-evoked currents in terms of both the amplitude and shift in the reversal potential, which signifies the existence of Na^+^ selective currents (Fig. 6f-h). These observations are in good agreement with the results for Glu447 and Phe514 in XtTPC3, which highlights the similarities between the two distinct gating mechanisms defined by the DII-S4 conformation shared in TPC subtypes.

### Naringenin has opposite effects on desipramine/voltage- and PI(3, 5)P_2_-dependent modes in HsTPC2

Naringenin is a plant flavonoid that exhibits a variety of biological activities and has been reported to inhibit the HsTPC2 current induced by PI(3, 5)P_2_ (41). In our experiments using the *Xenopus* oocyte expression system, although the results were not statistically significant, naringenin showed a tendency to suppress the currents induced by PI(3, 5)P_2_ intracellular injection, which is consistent with a previous report (Fig. 7a, b). On the other hand, naringenin unexpectedly showed potentiation of desipramine-induced currents and a shift of the G-V relationship toward a negative direction (Fig. 7c-f, Supplementary Table 1). Naringenin alone had no effect on HsTPC2 basal currents (Fig. 7d, e). Taken together, it is likely that naringenin is not a simple inhibitor of HsTPC2 but rather a so-called “biased modulator” that changes the equilibrium between the two modes in favor of the LyNaVA mode. These complex pharmacological effects of naringenin demonstrate the existence of a multimodal gating mechanism in TPC2 and provide new insights into the function and pharmacology in TPCs.

**Figure 7.**
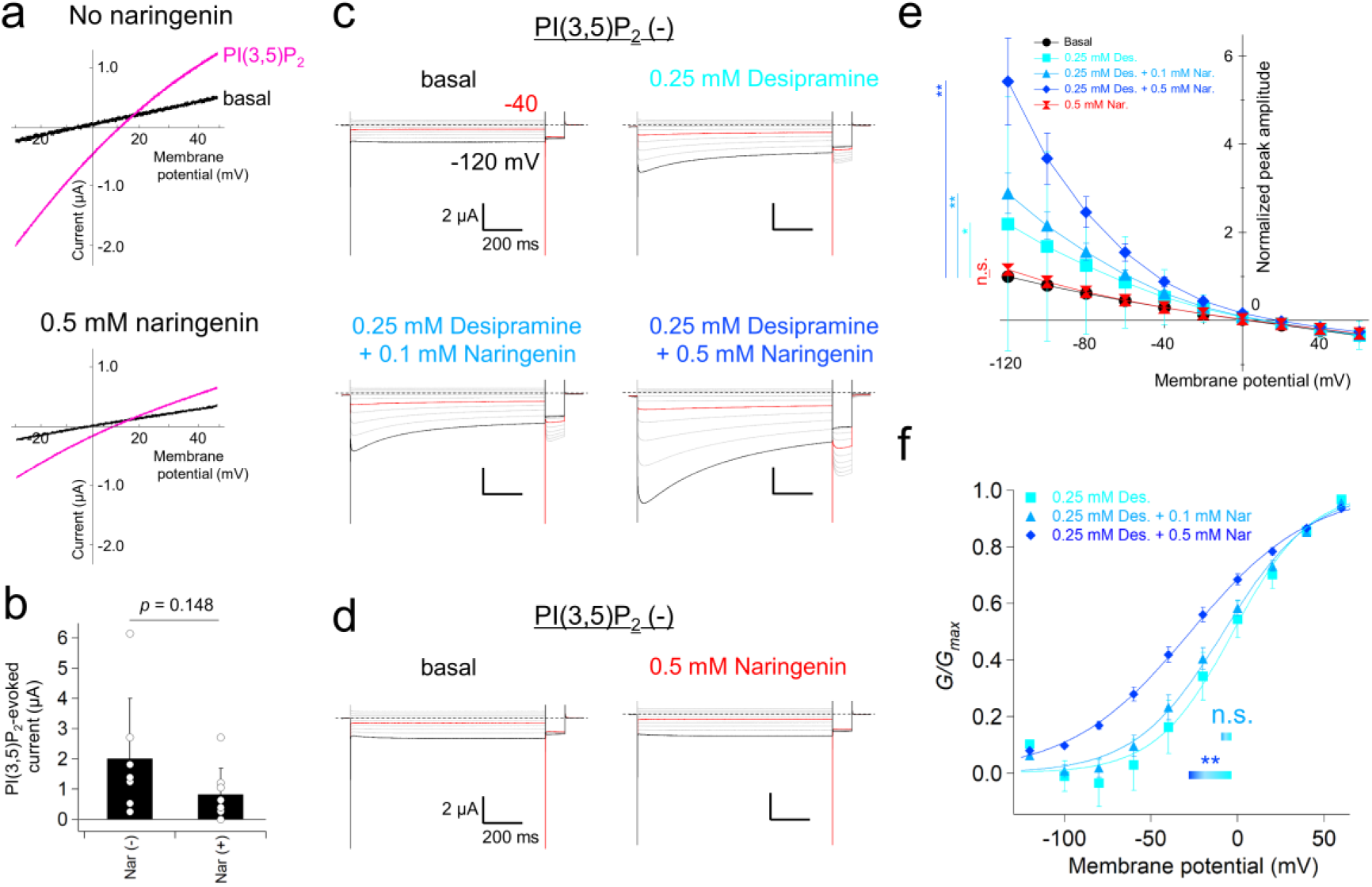
Different effects of naringenin on PI(3,5)P_2_-evoked and desipramine-induced currents in HsTPC2. (a) Representative recordings of PI(3, 5)P_2_-evoked currents of WT (ΔN37) HsTPC2 in the absence (top) and the presence (bottom) of 0.5 mM naringenin in the bath solution. Currents were evoked by ramp pulses ranging from -30 mV to +50 mV. (b) Plot for the current amplitude evoked by PI(3, 5)P_2_ obtained in (a) (n = 7-8). (c) Representative current recordings induced by desipramine and by the following co-application of desipramine and naringenin. Currents were elicited by step pulses, ranging from -120 mV to +60 mV, from a holding potential at 0 mV, followed by a -100 mV pulse to record the tail current. (d) Representative current recordings before (left) and after (right) the application of 0.5 mM naringenin. Currents were elicited by the same pulse protocol used in (c). (e) Normalized peak current amplitude at each membrane potential, obtained from the currents shown in (c) and (d) (n = 4-5). The peak currents were normalized to those in the absence of reagents in each recording. (f) G-V relationships for the currents induced by desipramine alone or by the co-application of desipramine and naringenin (n = 5). Statistical significance was examined by unpaired t-test or One-way ANOVA followed by Dunnett’s test in (b) and (e, f), respectively. **, * and n.s. mean *p* < 0.01, *p* < 0.05 and p > 0.05, respectively.

## Discussion

In this study, we revealed that the VSD2 conformations defines the complex switching and integrating mechanisms for PI(3, 4)P_2_- and voltage-gating in XtTPC3. We also confirmed the similarities between these mechanisms in XtTPC3 and LyNaVA- induced voltage gating and PI(3, 5)P_2_-gating in HsTPC2. The notion of the two distinct modes in HsTPC2 is supported by the finding that naringenin is a unique effector that changes the equilibrium between the two modes.

Our VCF analysis in XtTPC3 revealed peculiar relationships between the transfer of gating charges and the two steps of the state transitions (Fig. 5e). The F2 movement induces a transition from the intermediate state, which can open with a low open probability (I/O state), to the fully activated state. The A/O state likely plays a role in increasing the open probability more so than the I/O state. The F2 movement occurs along with most of the gating charge transfer (30), which is consistent with its large movement, where Phe514 on DII-S4 becomes closer to Glu438 (Fig. 2e). In the absence of PI(3, 4)P_2_, the F2 component and current occurred almost simultaneously (Fig. 5c-d). This coincidence between the current and fluorescence, which tracks the movement of the gating charge, is occasionally observed in some voltage-dependent ion channels, including KCNQ1(42). This means that owing to the poor coupling between the pore gate and each VSD, a single VSD movement can lead to gate opening (43). Therefore, F2 movement is still typical compared to S4 movement in other voltage-gated cation channels. On the other hand, the F1 component is quite atypical in that it can induce channel opening even before a large transfer of gating charges. A very small transfer of gating charges in the F1 component indicates a quite limited and local movement of DII-S4 (30). Consistent with this, VCF measurement with the fluorophore attached to the cytosolic end of DII-S4 in XtTPC3 showed no F1-corresponding movement (30). In addition, the gentle voltage dependence of the F1 component suggests that the intermediate conformation of VSD2 is similar to, and easily returns to, the resting conformation (Fig. 1e). This instability could explain why the I/O state was quite minor in WT XtTPC3 (Fig. 8a). An artificial interaction between Glu447 and Phe514, which stabilizes the intermediate conformation, increases the chance of the channels opening by reaching the I/O state. Interestingly, a recent report of the cryo-EM structures of plant TPC1, which does not have any PIP_2_ sensitivity, showed the conformational rearrangement of DII-S4 that is limited around the extracellular end by comparing the two “S4-down” conformations (44). This rearrangement induced a slight expansion of the entire VSD2 structure, as if it released DII-S4 for further movement. The F1 movement may correspond to this local structural rearrangement to prepare for the subsequent voltage-dependent F2 movement as well as for PIP_2_-gating (Fig. 8a). These observations indicate the multiple-state transition mechanism achieved by atypical F1 and typical F2 movements of DII-S4 in TPCs.

**Figure 8.**
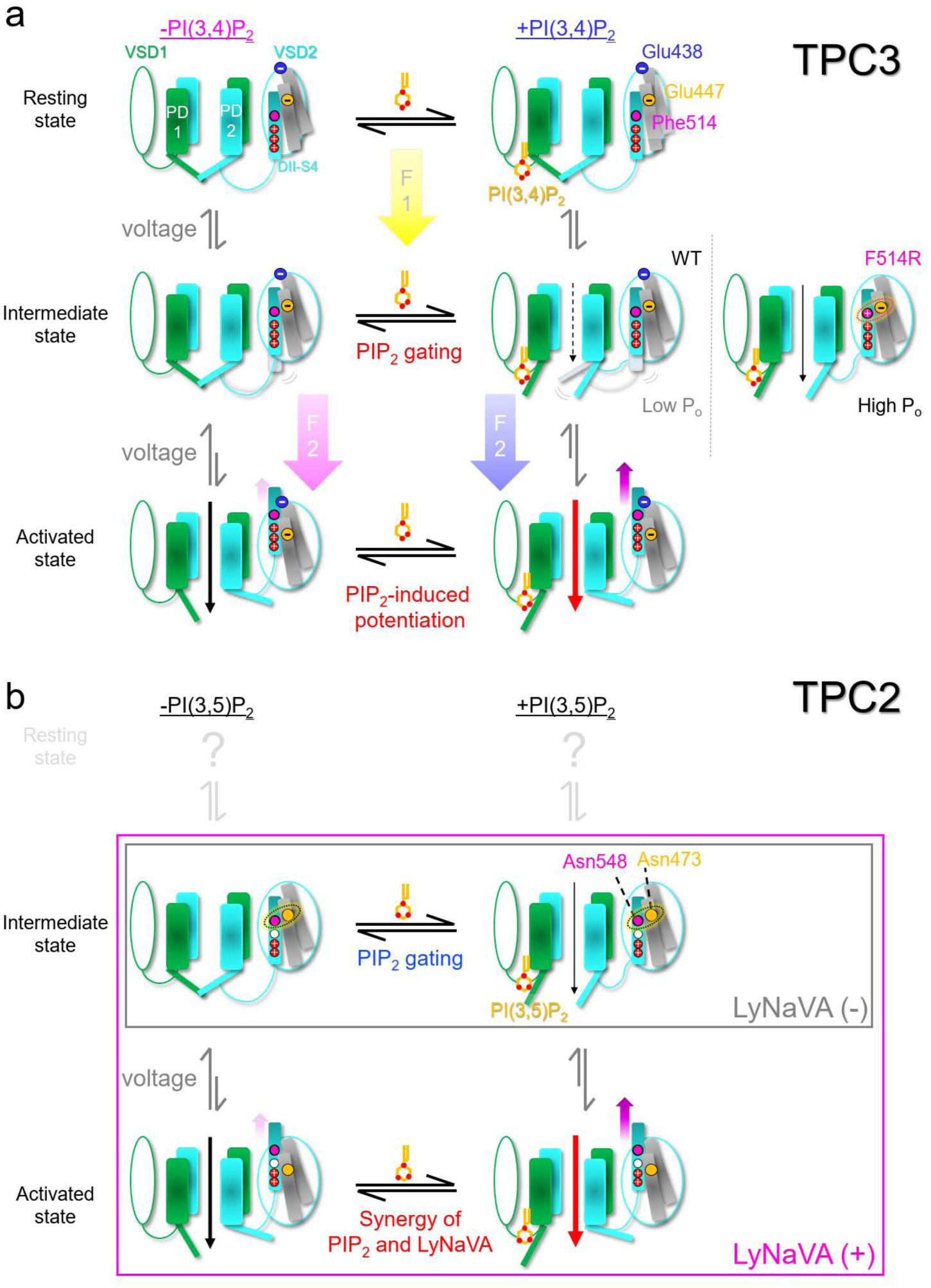
Schematic models for the state transitions in TPC3 and TPC2. (a) State transitions of TPC3 in the absence (left) and presence (middle) of PI(3, 4)P_2_. Voltage-dependent transition is shown in a vertical direction from resting (top), to intermediate (middle) and activated states (bottom). Large arrowheads between the states indicate the F1 (yellow) and F2 (magenta and blue) components in the VCF measurement, respectively. In the intermediate state of WT (middle), the activation gate could open with a low open probability in the presence of PI(3, 4)P_2_, possibly due to the instability of this conformation. In the F514R mutant (middle right), the electrostatic interaction between Glu447 and F514R (red dotted oval) stabilizes the conformation, resulting in a high open probability. In the activated state, the open probability in the presence of PI(3, 4)P_2_ is much higher than in its absence, as PI(3, 4)P_2_ binding facilitates the DII-S4 movement through the inter-domain interaction between DI-S6 and DII-S6. (b) State transitions of TPC2, similar to (a). In TPC2, it is unclear whether the states corresponding to the resting states of TPC3 exists or not. A hydrogen bond between Asn473 and Asn548 (dotted oval) stabilizes the PIP_2_-preferable intermediate state (middle). Voltage-dependent state transitions from the intermediate states to the activated states are invisible in the absence of LyNaVA, and thus only the PIP_2_-dependent gating occurs. LyNaVA binding allows the currents by the voltage-dependent transition. Further increase in the open probability occurs when PI(3, 5)P_2_ and voltage are given simultaneously(33), which corresponds to the PI(3, 4)P_2_-induced potentiation of voltage dependence of TPC3 shown in (a).

The unique PIP_2_-dependence revealed in XtTPC3 could be a new type of PIP_2_- dependent regulatory mechanism in voltage-gated ion channels. A PI(4, 5)P_2_ binding site in KCNQ1 is located at the interface between the S4-S5 linker and VSD (45, 46), similar to those of PI(3, 4)P_2_ and PI(3, 5)P_2_ in TPCs that are near their DI-S4/S5 linkers (26, 31, 32, 37). The binding of PI(4, 5)P_2_ to KCNQ1 plays a pivotal role in the coupling of VSDs and PDs, and its dissociation results in a remarkable decrease in channel activity (46). Because it occurs even if KCNQ1 is co-expressed with its auxiliary subunit E1 or E3, which changes the AO/IO ratio, PI(4, 5)P_2_ binding is considered to be required regardless of the state (38, 47, 48). In the case of XtTPC3, analysis of the LV components of F514R and E447R/F514E indicated that the binding of PI(3, 4)P_2_ is indispensable for the I/O state (Fig. 2f and 4c). However, the loss of PI(3, 4)P_2_ shifted the voltage-dependence in the WT current and in the HV component of F514R, which mainly reflects the A/O state, toward the depolarized direction, but the channel currents were not lost. These observations demonstate a unique PIP_2_-dependent regulatory mechanism that focuses more specifically on the intermediate state than on the activated state.

Because the I/O state in WT XtTPC3 in the presence of PI(3, 4)P_2_ adds a small conductance at the hyperpolarized potentials where F1 is dominant (Fig. 1c), it would be the cause for the apparent shift in the voltage dependence toward the negative direction, together with the A/O conductance in the depolarized voltage range. In addition, the potentiation mechanism may be, at least partly, due to the easier transition from the I/C state to the A/O state through the I/O state, rather than through the A/C state (Fig.5e). The voltage dependence of F2, which reflects the transition from the intermediate to activated states, is shifted toward the negative direction in the presence of PI(3, 4)P_2_ compared to in its absence (Fig. 5b and Hirazawa et al. (30)). The I/O state takes the open conformation of the activation gate, which is composed of DI-S6 and DII-S6. Because DII-S4 becomes more movable upon PI(3, 4)P_2_ binding likely through the inter-domain interaction between DI-S6 and DII-S6 and the coupling between DII-S4 and DII-S6 (30, 31), the I/O state may give DII-S4 a better chance to reach the fully activated conformation compared to the I/C state, where the DII-S6 would remain closed through the PI(3, 4)P_2_-unbound DI-S6 (Fig. 8a). These mechanisms could be the molecular basis for the synergy between the two domains receiving distinct stimuli of voltage and PIP_2_.

In this study, DII-S4 in HsTPC2 was found to play a major role in LyNaVA- induced voltage dependence. The voltage sensitivity of DII-S4 in TPC2 was not impaired, but it might be decoupled with the activation gate. The binding of LyNaVAs is thought to restore this coupling, through an unknown mechanism. The synergy between LyNaVAs and PI(3, 5)P_2_ in HsTPC2, which has been shown in a previous report (33), appears to be homologous to the PI(3, 4)P_2_-dependent potentiation of voltage-dependence in XtTPC3. This homology strongly suggests shared and fundamental activation mechanisms in the TPC family (Fig. 8). A hydrogen bond between Asn473 and Asn548 observed in the cryo-EM structure of HsTPC2 suggests that it was originally tuned to be locked in the intermediate state to allow PIP_2_-gating (Fig. 2c, 8b). Furthermore, a comparison between HsTPC2 and the XtTPC3 model, as well as in a mouse TPC1 (MmTPC1), showed that the relative positions of DII-S4 within VSD2 were almost identical and possibly captured in the intermediate state (Supplementary Fig. 6), indicating that this conformation is homologous and stable in all three subtypes. TPC subtypes are thought to share the fundamental framework enabling both PIP_2_-gating and voltage gating, and achieve optimal gating by changing the mode and extent they suppress. TPC2 is biased to PIP_2_-gating, while TPC3 is to voltage gating. Agents such as LyNaVAs, whose binding could induce conformational rearrangements that release these suppressions, may have the potential to act as TPC agonists (Fig. 8b). In summary, TPC subtypes might share gating mechanisms and be tuned to achieve their optimal extent of balanced gating for voltage and PIP_2_.

The different effects of naringenin on PI(3, 5)P_2_-evoked current and desipramine-induced currents suggest that they are distinct activation mechanisms, proving the existence of multimodal gating mechanisms in HsTPC2 (Fig. 7). Naringenin inhibited PI(3, 5)P_2_-induced TPC2 currents (41) and prevented the intracellular entry of several respiratory infectious viruses, including SARS-CoV-2, in *in vitro* experiments (17). The effects of naringenin revealed in this study provide interesting insights into strategies for the development of TPC2-targeted drugs. Although there is currently no evidence that the LyNaVA mode functions under physiological conditions, its distinct gating mechanism from the PIP_2_-mode raises the possibility that it potentially reflects a physiological role under specific conditions. This notion seems to be analogous to a report stating that TPC2 currents with different ion selectivity, which were induced by the two distinctive agonists, have different physiological consequences in the lysosomes (34). It is also possible that the inhibitory effect of naringenin on the intracellular entry of SARS-CoV-2 is caused not only by inhibition of the PIP_2_ current but also by induction of the LyNaVA-mode. Given these findings, the mode-targeted pharmacology of TPC2, with naringenin-like “biased modulators,” might be interesting and worth considering as a target to prevent the invasion of diverse viruses.

In summary, the present study showed that VSD2 in TPCs plays an active role in integrating or switching the PIP_2_-gating and voltage-gating modes, which is an unprecedented mechanism among voltage-gated ion channels. The intermediate conformation is defined as the PIP_2_-dependent mode. The balance between two modes is likely to be tuned in each TPC subtype and could be switched or biased by changing the VSD2 conformation, or especially in TPC2 by a unique type of “biased modulator”. Their multimodality and regulation may be a direction for drug development to target TPCs.

## Material and Methods

### Ethical approval

All animal experiments were approved by the Animal Care Committee of the National Institutes of Natural Sciences (an umbrella institution of the National Institute for Physiological Sciences, Japan), and performed following its guidelines.

### cRNA preparation and injection to *Xenopus* oocytes

Preparation and injection of cRNAs were performed as previously described (31). The cDNAs of HsTPC2, HsIP4B, and HsIP5D were purchased from Genscript and subcloned into pGEMHE. The cRNAs were synthesized with mMESSAGE mMACHINE T7 kit (Thermo Fisher Scientific) from the linearized cDNAs. The oocytes were collected from *Xenopus laevis* (Hamamatsu Seibutsu Kyouzai). Frogs were anesthetized by 0.15% tricaine, and oocytes were surgically removed as previously described(49). Then, the oocytes were kept at 17 ⁰C in frog Ringer’s solution containing (in mM): 88 NaCl, 1 KCl, 2.4 NaHCO_3_, 0.3 Ca(NO_3_)_2_, 0.41 CaCl_2_ and 0.82 MgSO_4_, 15 Hepes (pH7.6) with 0.1% penicillin–streptomycin (Sigma-Aldrich). The oocytes were injected with 50 nl of 0.5 mg/ml cRNA solution and incubated at 17 ⁰C in frog Ringer’s solution. In the case of co-expression, 16.7 ng of XtTPC3 cRNA and 8.3 ng of HsIP4B or HsIP5D cRNA were mixed and injected. Currents were measured 2 to 4 days after injection, depending on the current amplitude.

### Electrophysiological recording of oocytes

Currents were measured under two-electrode voltage clamp mode using an OC- 725C amplifier (Warner Instruments) and pClamp10.7 software (Molecular Devices). Data were digitized at 10 kHz through Digidata1440 (Molecular Devices). The resistance of microelectrodes was adjusted to be 0.2–0.5MΩ when filled with the solution of 3M K-acetate and 10mM KCl. The recording bath solution is, unless specially described in each legend, ND-96, which contains (in mM): 96 NaCl, 2 KCl, 1.8 CaCl_2_, 1 MgCl_2_ and 5 HEPES-NaOH (pH 7.4). To avoid the damage derived from the sustained depolarization in XtTPC3 or HsTPC2, the oocytes expressing the channels were kept in the ND96-based solutions with low or medium Na^+^ concentration, where 96 mM NaCl was replaced with 5 mM NaCl and 91 mM N-Methyl D-glucamine (NMDG) (low) or with 20 mM NaCl and 76 mM NMDG-HCl (medium). The recordings in XtTPC3 co-expressed with HsIP5D were performed in the ND-96 solution containing 10 µM insulin (Wako), which further increases the PI(3, 4)P_2_ concentration through a signaling pathway endogenous in *Xenopus* oocytes (50). The recordings from XtTPC3 co-expressed with HsIP4B require sufficient time intervals of more than 10 sec every traces, to return to the PI(3, 4)P_2_-depleted condition. It is because strong depolarizing pulses activate the voltage-dependent PI(3, 4)P_2_ producing activity endogenous in *Xenopus* oocytes and instantaneously lead to the partial potentiation of XtTPC3 (31). In the measurement of desipramine-induced HsTPC2 currents, the pre- pulses at +50 mV for 2 sec were applied in each trace to prevent the run-down by an unknown mechanism.

In PI(3, 5)P_2_-injection experiments, 5 mM PI(3, 5)P_2_–diC8 (Echelon Biosciences) was manually injected by positive pressure using the glass needle filled with the PI(3, 5)P_2_ solution. The recordings for PI(3, 5)P_2_ injection were performed under ND-96 solution containing 10 µM Ani9 (Sigma-Aldrich) to inhibit the endogenous Ca^2+^- activated Cl^-^ channel, which could be weakly activated by PI(3, 5)P_2_. Cu-Phen solution used for double cysteine mutants contained 2 µM CuSO_4_ (Wako) and 10 µM phenanthroline (Wako) which were diluted to ND-96 immediately before each experiment. The oocytes expressing some double cysteine mutants were required to be pre-treated by 0.5 mM DTT (Wako) for more than 30 min in low or medium Na^+^ solution to make their channel activity recovered. Riluzole and naringenin were purchased from Tokyo Chemical Industry, and desipramine was from Sigma.

### Voltage clamp fluorometry

The oocytes expressing Q507C or E447R/Q507C/F514E XtTPC3 were labeled with 100 µM Alexa Fluor™ 488 C5 Maleimide (Thermo Fisher Scientific) at 25 °C for 1.5 hours in the ND-96 based NMDG solution which contains (in mM) 98 NMDG, 1.8 CaCl_2_, 1 MgCl_2_, and 5 HEPES-NMDG, pH 7.4. The oocytes were then rinsed twice and kept in the NMDG solution.

The recordings were performed with an upright fluorescence microscope (Olympus BX51WI) equipped with a water immersion objective lens (Olympus XLUMPLFLN 20x/1.00). The excitation light from the xenon arc lamp (Hamamatsu Photonics) was applied through a band-pass excitation filter (470-490 nm). The intensity of the excitation light was decreased to 1.5 or 6.0% with ND filters (U-25ND6, U-25ND25 Olympus) to avoid bleaching induced by too strong illumination. The emission light was passed through the 510 -550 nm band pass filter (U-MNIBA2, Olympus), and introduced to a photomultiplier (Hamamatsu Photonics). The fluorescent signal and current were acquired by the Digidata 1332 (Axon Instruments) and pClamp 10.3 software (Molecular devices) at 100 kHz.

The data were collected 5 times and averaged to improve the signal noise ratio. A slow bleaching was adjusted by assuming that the decrease until the first step pulse was linear. Due to the large current amplitude needed to collect sufficient fluorescence in VCF experiments, it was inevitable that voltages for the tail currents were occasionally not well clamped, especially in E447R/Q507C/F514E whose expression strongly impaired oocyte condition. We set less than 6 mV volage-error on the maximum step pulse of +200 mV from a holding potential at -60 mV, as an acceptable range, demanding cautions in the accuracy of the data at high voltages.

### Data analysis and statistics

Electrophysiological data were analyzed using Clampfit 10.7 (Molecular Devices) and Igor Pro (WaveMetrics). Tail currents, elicited by the constant voltage after the varying step pulses, were used to obtain G–V relationships. The voltage for tail currents was + 60 mV unless specially described in the figures or in the figure legends. The G–V relationships were calculated by fitting to a two-state Boltzmann equation as follows;

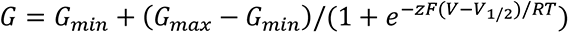

G_min_ was set to zero and then *G*/*G*_max_ was plotted against membrane voltage.

Time constants for activation in current and fluorescence were obtained using Clampfit by fitting each trace with a single exponential function. A range of 100 ms after step pulses was used for fitting in both the fluorescent and current traces because the following decrease in currents after 100 ms made fitting and interpretation complicated. Data are presented as mean ± S.D. Student’s *t*-test or one-way ANOVA, followed by Dunnett’s test, were used depending on the experiments, to examine statistical significance. *p* < 0.05 was considered statistically significant.

### Modeling of the XtTPC3 structure

A homology model of XtTPC3 was generated based on the cryo-EM structure of *Danio rerio* TPC3 (PDB: 6V1Q) using SWISS-MODEL (51). The structures of PI(3, 5)P_2_-bound MmTPC1 and PI(3, 5)P_2_-bound open HsTPC2 (PDB: 6c9a and 6nq0, respectively) were used for structural comparison with the XtTPC3 model. All the figures were made with PyMol (Schrödinger).

### Data availability

All the data supporting this study are available within this article and supplementary data.

## Acknowledgment

We thank Ms. Chizue Naito and Ms. Tomomi Yamamoto for technical assistance and all members of Kubo laboratory for discussion. This study was supported by the Hiroshi and Aya Irisawa Memorial Promotion Award for Young Physiologists (to T.S.) from the Physiological Society of Japan, and the Grants-in-Aid (C) 20K07284 (to T.S.), (B) 17H04021 (to Y.K.) and (B) 20H03424 (to Y.K.) from Japan Society for the Promotion of Science.

## Author contributions

T.S. and Y.K. conceived the experiment. T.S. collected and analyzed the electrophysiological data. K.H. performed the optimization of experimental conditions. T.S. wrote the draft of the manuscript. T.S., K.H., and Y.K. completed the manuscript. All authors approved the final version of the manuscript.

## References

1. P. J. Calcraft, et al., NAADP mobilizes calcium from acidic organelles through two-pore channels. Nature 459, 596–600 (2009).

2. E. Brailoiu, et al., Essential requirement for two-pore channel 1 in NAADP-mediated calcium signaling. J. Cell Biol. 186, 201–209 (2009).

3. X. Zong, et al., The two-pore channel TPCN2 mediates NAADP-dependent Ca(2+)-release from lysosomal stores. Pflugers Arch. 458, 891–9 (2009).

4. H. Xu, D. Ren, Lysosomal Physiology. Annu. Rev. Physiol. 77, 57–80 (2015).

5. Q. Zhang, et al., Dynamic PIP2 interactions with voltage sensor elements contribute to KCNQ2 channel gating. Proc Natl Acad Sci U S A 110, 20093– 20098 (2013).

6. A. Faviaa, et al., VEGF-induced neoangiogenesis is mediated by NAADP and two-pore channel-2 -dependent ca2+signaling. Proc. Natl. Acad. Sci. U. S. A. 111, 4706–4715 (2014).

7. Y. K. Chao, et al., TPC2 polymorphisms associated with a hair pigmentation phenotype in humans result in gain of channel function by independent mechanisms. Proc. Natl. Acad. Sci. U. S. A. 114, E8595–E8602 (2017).

8. C. Grimm, et al., High susceptibility to fatty liver disease in two-pore channel 2- deficient mice. Nat. Commun. 5 (2014).

9. L. N. Hockey, et al., Dysregulation of lysosomal morphology by pathogenic LRRK2 is corrected by TPC2 inhibition. J. Cell Sci. 128, 232–238 (2015).

10. Y. Sakurai, et al., Two-pore channels control Ebola virus host cell entry and are drug targets for disease treatment. Science 347, 995–998 (2015).

11. B. Hu, H. Guo, P. Zhou, Z. L. Shi, Characteristics of SARS-CoV-2 and COVID- 19. Nat. Rev. Microbiol. 19, 141–154 (2021).

12. M. Blaess, L. Kaiser, M. Sauer, R. Csuk, H. P. Deigner, COVID-19/SARS-CoV- 2 infection: Lysosomes and lysosomotropism implicate new treatment strategies and personal risks. Int. J. Mol. Sci. 21, 1–29 (2020).

13. G. S. Gunaratne, J. S. Marchant, The ins and outs of virus trafficking through acidic Ca2+ stores. Cell Calcium 102 (2022).

14. X. Ou, et al., Characterization of spike glycoprotein of SARS-CoV-2 on virus entry and its immune cross-reactivity with SARS-CoV. Nat. Commun. 11 (2020).

15. L. Riva, et al., Discovery of SARS-CoV-2 antiviral drugs through large-scale compound repurposing. Nature 586, 113–119 (2020).

16. Y. L. Kang, et al., Inhibition of PIKfyve kinase prevents infection by Zaire ebolavirus and SARS-CoV-2. Proc. Natl. Acad. Sci. U. S. A. 117, 20803–20813 (2020).

17. N. Clementi, et al., Naringenin is a powerful inhibitor of SARS-CoV-2 infection in vitro. Pharmacol. Res. 163 (2021).

18. E. Peiter, et al., The vacuolar Ca2+-activated channel TPC1 regulates germination and stomatal movement. Nature 434, 404–8 (2005).

19. F. H. Yu, W. A. Catterall, The VGL-chanome: a protein superfamily specialized for electrical signaling and ionic homeostasis. Sci. STKE 2004, re15 (2004).

20. J. Guo, et al., Structure of the voltage-gated two-pore channel TPC1 from Arabidopsis thaliana. Nature 531, 196–201 (2016).

21. A. F. Kintzer, R. M. Stroud, Structure, inhibition and regulation of two-pore channel TPC1 from Arabidopsis thaliana. Nature 531, 258–264 (2016).

22. C. M. Armstrong, F. Bezanilla, W. Hole, Charge movement associated with the opening and closing of the activation gates of the Na channels. J. Gen. Physiol. 63, 533–52 (1974).

23. S. K. Aggarwal, R. MacKinnon, Contribution of the S4 segment to gating charge in the Shaker K+channel. Neuron 16, 1169–1177 (1996).

24. S. A. Seoh, D. Sigg, D. M. Papazian, F. Bezanilla, Voltage-sensing residues in the S2 and S4 segments of the Shaker K+channel. Neuron 16, 1159–1167 (1996).

25. M. S. Dickinson, A. Myasnikov, J. Eriksen, N. Poweleit, R. M. Stroud, Resting state structure of the hyperdepolarization activated two-pore channel 3. Proc. Natl. Acad. Sci. U. S. A. 117, 1988–1993 (2020).

26. J. She, et al., Structural insights into the voltage and phospholipid activation of the mammalian TPC1 channel. Nature 556, 130–134 (2018).

27. X. Wang, et al., TPC proteins are phosphoinositide- Activated sodium-selective ion channels in endosomes and lysosomes. Cell 151, 372–383 (2012).

28. C. Cang, B. Bekele, D. Ren, The voltage-gated sodium channel TPC1 confers endolysosomal excitability. Nat. Chem. Biol. 10, 1–8 (2014).

29. C. Cang, et al., mTOR regulates lysosomal ATP-sensitive two-pore Na(+) channels to adapt to metabolic state. Cell 152, 778–90 (2013).

30. K. Hirazawa, M. Tateyama, Y. Kubo, T. Shimomura, Phosphoinositide regulates dynamic movement of the S4 voltage sensor in the 2nd repeat in Two-pore channel 3. J. Biol. Chem. 297, 101425 (2021).

31. T. Shimomura, Y. Kubo, Phosphoinositides modulate the voltage dependence of two-pore channel 3. J. Gen. Physiol. 151, 986–1006 (2019).

32. J. She, et al., Structural mechanisms of phospholipid activation of the human TPC2 channel. Elife 8, 1–17 (2019).

33. X. Zhang, et al., Agonist-specific voltage-dependent gating of lysosomal two- pore na+ channels. Elife 8, 1–18 (2019).

34. S. Gerndt, et al., Agonist-mediated switching of ion selectivity in TPC2 differentially promotes lysosomal function. Elife 9, 1–63 (2020).

35. L. M. Mannuzzu, M. M. Moronne, E. Y. Isacoff, Direct physical measure of conformational rearrangement underlying potassium channel gating. Science 271, 213–6 (1996).

36. T. Balla, Phosphoinositides: Tiny Lipids With Giant Impact on Cell Regulation. Physiol. Rev. 93, 1019–1137 (2013).

37. M. S. Dickinson, A. Myasnikov, J. Eriksen, N. Poweleit, R. M. Stroud, Resting state structure of the hyperdepolarization activated two-pore channel 3. Proc. Natl. Acad. Sci. U. S. A. 117, 1988–1993 (2020).

38. K. C. Taylor, et al., Structure and physiological function of the human KCNQ1 channel voltage sensor intermediate state. Elife 9 (2020).

39. E. Brailoiu, et al., An NAADP-gated two-pore channel targeted to the plasma membrane uncouples triggering from amplifying Ca2+ signals. J. Biol. Chem. 285, 38511–6 (2010).

40. Y. Lin-Moshier, et al., The Two-pore channel (TPC) interactome unmasks isoform-specific roles for TPCs in endolysosomal morphology and cell pigmentation. Proc. Natl. Acad. Sci. 111, 13087–13092 (2014).

41. I. Pafumi, et al., Naringenin Impairs Two-Pore Channel 2 Activity And Inhibits VEGF-Induced Angiogenesis. Sci. Rep. 7 (2017).

42. J. D. Osteen, et al., KCNE1 alters the voltage sensor movements necessary to open the KCNQ1 channel gate. Proc. Natl. Acad. Sci. U. S. A. 107, 22710–22715 (2010).

43. J. Cui, Voltage-Dependent Gating: Novel Insights from KCNQ1 Channels. Biophys. J. 110, 14–25 (2016).

44. M. S. Dickinson, et al., Molecular basis of multistep voltage activation in plant two-pore channel 1. Proc. Natl. Acad. Sci. U. S. A. 119 (2022).

45. J. Sun, R. MacKinnon, Structural Basis of Human KCNQ1 Modulation and Gating. Cell 180, 340–347.e9 (2020).

46. M. A. Zaydman, et al., Kv7.1 ion channels require a lipid to couple voltage sensing to pore opening. Proc. Natl. Acad. Sci. U. S. A. 110, 13180–5 (2013).

47. R. Barro-Soria, M. E. Perez, H. P. Larsson, KCNE3 acts by promoting voltage sensor activation in KCNQ1. Proc. Natl. Acad. Sci. U. S. A. 112, E7286–E7292 (2015).

48. A. A. Royal, A. Tinker, S. C. Harmer, Phosphatidylinositol-4,5-bisphosphate is required for KCNQ1/KCNE1 channel function but not anterograde trafficking. PLoS One 12 (2017).

49. I. S. Chen, et al., Non-sedating antihistamines block G-protein-gated inwardly rectifying K+ channels. Br. J. Pharmacol. 176, 3161–3179 (2019).

50. L. Liu, et al., A glutamate switch controls voltage-sensitive phosphatase function. Nat. Struct. Mol. Biol. 19, 633–41 (2012).

51. T. Schwede, J. Kopp, N. Guex, M. C. Peitsch, SWISS-MODEL: An automated protein homology-modeling server. Nucleic Acids Res. 31, 3381–3385 (2003).

