## Supplementary figures and table for "Conformational rearrangements in 2^nd^ voltage sensor domain switch PIP_2_- and voltage-gating modes in two-pore channels"

**This PDF file includes:**

Supplementary text
Figures S1 to S6
Tables S1

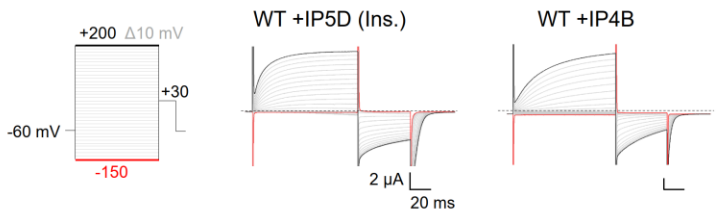


**Fig. S1.** Representative current recordings of XtTPC3 WT, co-expressed with HsIP5D and treated by insulin (middle) and with HsIP4B (right). Currents were elicited by the pulse protocol shown on the left.

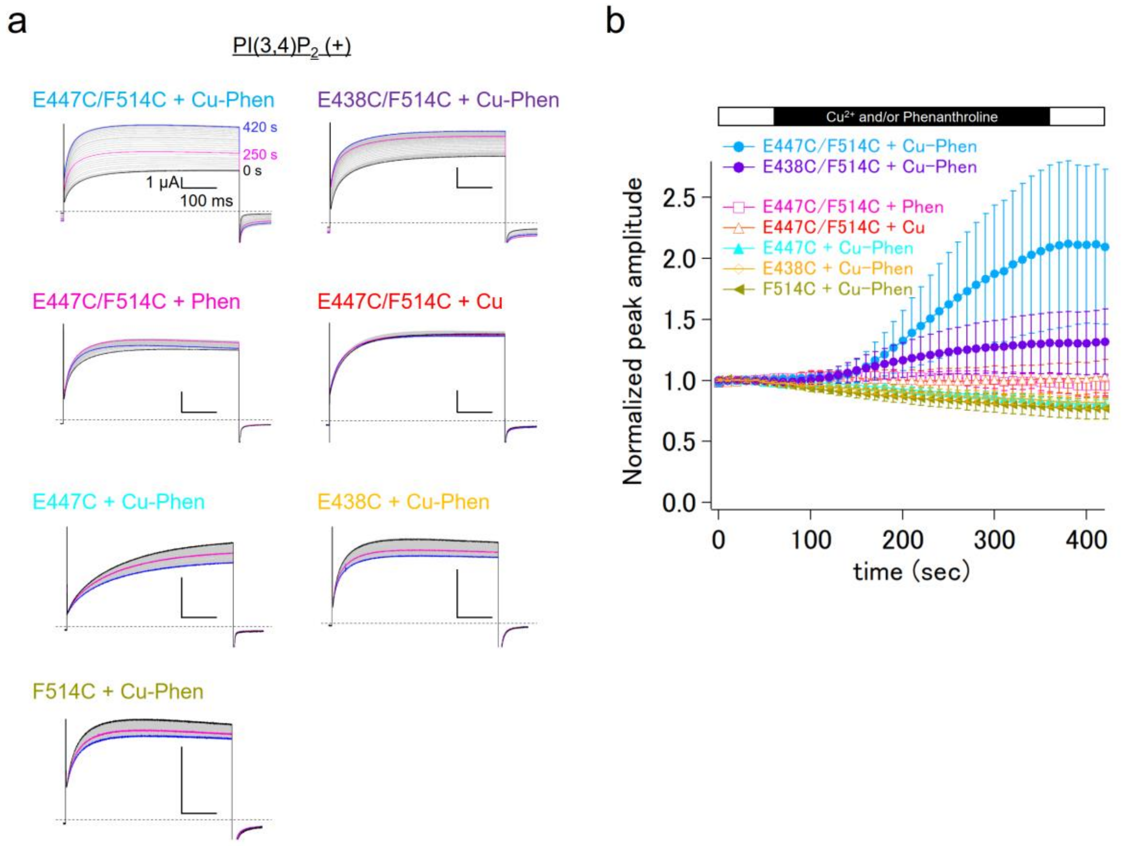


**Fig. S2.** (a) Representative current recordings for the effects of 2 μM $Cu^{2+}$ and/or 10 μM phenanthroline on a series of mutants. Currents were elicited by +120 mV step pulses from a holding potential at -60 mV in the ND96-based 20 mM $Na^{+}$ solutions. $Cu^{2+}$ and/or phenanthroline were perfused between 60 sec and 360 sec. (b) Time lapse changes of the normalized current amplitude are obtained from the data as shown in (a) (n = 3-8).

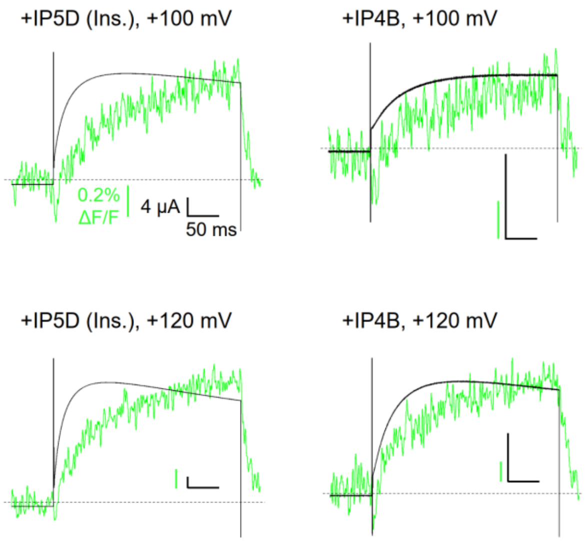


**Fig. S3.** Superimposed traces of current (black) and fluorescence (green) of Q507C co-expressed with HsIP5D and treated by insulin (left), and with HsIP4B (right) at +100 mV or +120 mV.

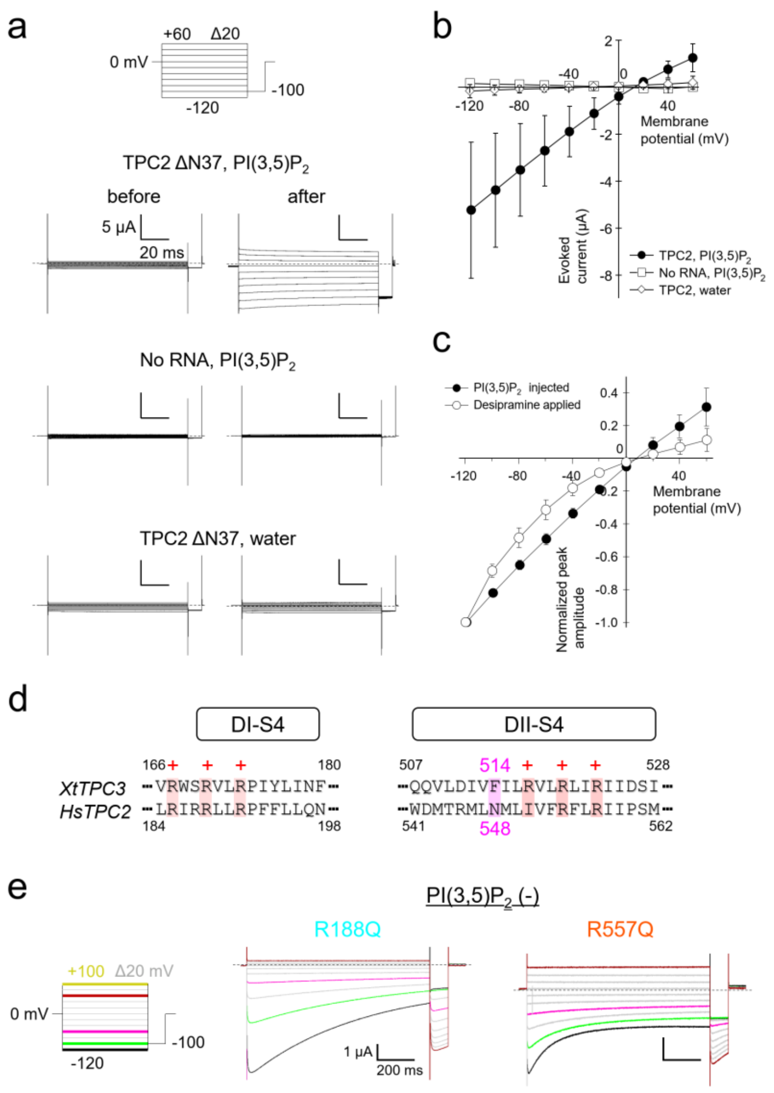


**Fig. S4.** (a) Representative current recordings from the oocytes expressing HsTPC2 ΔN 37 (WT) or with no cRNA injection, in response to the injection of PI(3,5)$P_2$ or water. The recordings from the same oocyte before (left) and after (right) injection are shown. (b) I-V relationships obtained from the peak current amplitude in (a) (n = 4). (c) Normalized peak current amplitude of PI(3,5)$P_2$-evoked currents and desipramine-induced currents (n = 4-12). The values immediately after the application of step pulses are plotted. (d) Amino acid sequence alignments for DI-S4 and DII-S4 in XtTPC3 and HsTPC2. The positions for the conserved positively charged amino acids are denoted as + at the top of their sequences. Phe514 in XtTPC3 and Asn548 in HsTPC2 are highlighted as magenta. (e) Representative current recordings for HsTPC2 R188Q and R557Q induced by 1 mM desipramine.

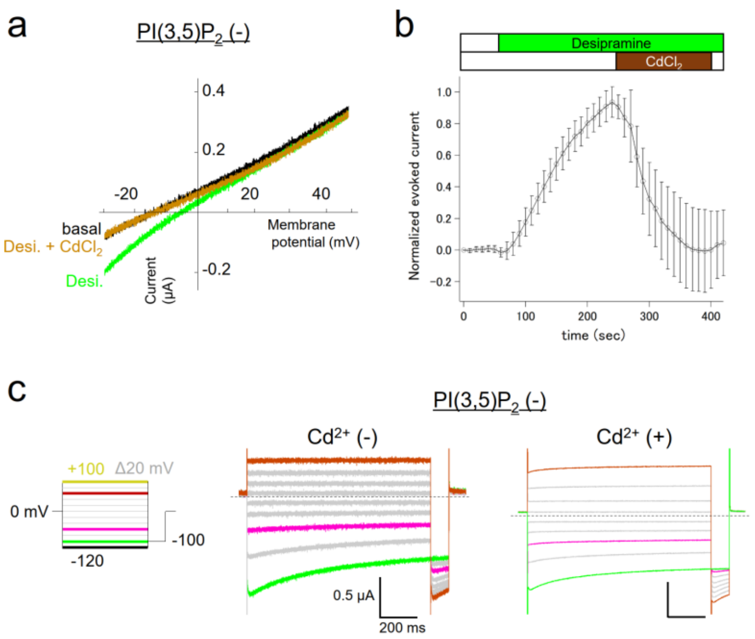


**Fig. S5.** (a) Representative current recordings evoked by repetitive ramp pulses of N473C/N548C, in response to the bath perfusion of 1 mM desipramine (brown) and the following application of 1 mM desipramine and 100 µM $CdCl_2$. (b) Time course of the current change obtained from the data in (a) (n = 6). The evoked current was normalized by the maximum evoked amplitude in the presence of 1 mM desipramine. (c) Representative current recordings of N473C/N548C induced by 1 mM desipramine in the absence (middle) or the presence (right) of 100 µM $CdCl_2$. Each recording was obtained from a different oocyte.

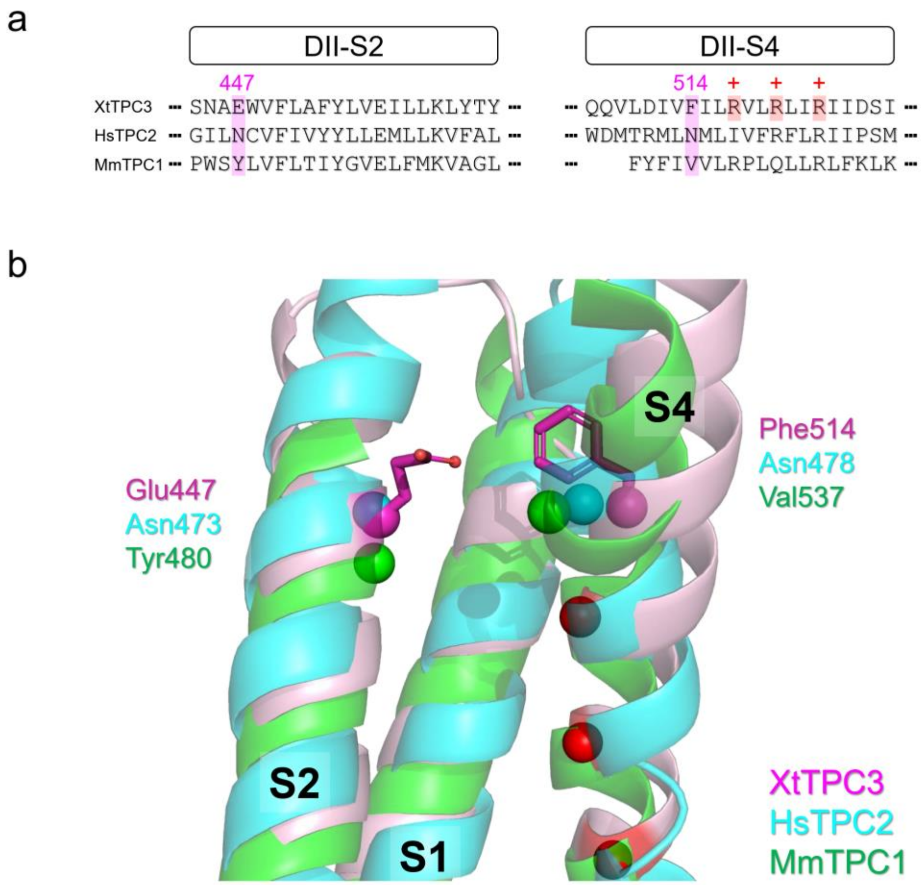


**Fig. S6.** (a) Sequence alignments for DII-S2 and DII-S4 in XtTPC3, HsTPC2 and MmTPC1. The positions for the positively charged amino acids in XtTPC3 are colored in red and denoted as + at the top of their sequences. The positions corresponding to Glu447 and Phe514 in XtTPC3 are highlighted as magenta. (b) The aligned VSD2 structures for XtTPC3 (light pink) HsTPC2 (aqua) and MmTPC1 (green). The Cα positions for the residues corresponding to Glu447 and Phe514 in XtTPC3 are shown as spheres in each color representing the belonging subtype. The side chains of Glu447 and Phe514 in XtTPC3 are shown as a stick model and colored in purple. Red spheres indicate the Cα positions of the positively charged arginines in XtTPC3.

| | | | | $V_{1/2}$ (mV) | *n* |
|---|---|---|---|---|---|
| | | Cu-Phen | | | |
| *XtTPC3* | | | | | |
| Q507C*Alexa-488 | +IP5D | | | 69.5 ± 0.84 | 6 |
| | +IP4B | | | 114 ± 0.46 | 8 |
| E447C/F514C | +IP5D | | | 144 ± 1.0 | 5 |
| | +IP5D | + | | 144 ± 1.1 | 5 |
| | +IP4B | | | 178 ± 3.2 | 5 |
| | +IP4B | + | | 173 ± 3.1 | 5 |
| E438C/F514C | +IP5D | | | 77.1 ± 0.83 | 7 |
| | +IP5D | + | | 78.4 ± 1.6 | 7 |
| | +IP4B | | | 137 ± 1.3 | 4 |
| | +IP4B | + | | 139 ± 1.6 | 4 |
| | Desipramine | Naringenin | $Cd^{2+}$ | | |
| *HsTPC2* | | | | | |
| WT (ΔN37) | 1 mM | | | -44.9 ± 4.2 | 6 |
| R185Q | 1 mM | | | -43.7 ± 1.6 | 4 |
| R188Q | 1 mM | | | -49.5 ± 0.79 | 4 |
| I551R | 1 mM | | | 80.5 ± 3.6 | 6 |
| R554Q | 1 mM | | | -6.00 ± 11 | 5 |
| R557Q | 1 mM | | | -11.0 ± 8.8 | 5 |
| N473C/N548C | 1 mM | | | -39.0 ± 10 | 7 |
| | 1 mM | | + | -12.6 ± 8.3 | 7 |
| N548C | 1 mM | | | -59.4 ± 6.7 | 5 |
| | 1 mM | | + | -52.0 ± 7.2 | 6 |
| WT | 0.25 mM | | | -2.88 ± 5.3 | 5 |
| | 0.25 mM | 0.1 mM | | -8.13 ± 3.4 | 5 |
| | 0.25 mM | 0.5 mM | | -27.1 ± 5.1 | 5 |

**Table S1.** The $V_{1/2}$ values in a series of mutants of XtTPC3 and HsTPC2. * indicates the labelling by Alexa Fluor-488 maleimide.
